# TCR-FramePose: a local-frame representation for decomposing global docking and CDR3 loop geometry in TCR-pMHC recognition

**DOI:** 10.64898/2026.06.30.735664

**Authors:** Kun Hee Kim, Xianli Jiang, Qing Ye, Vakul Mohanty, Merve Dede, Alexandre Reuben, Ken Chen

## Abstract

T cell receptor recognition of peptide–MHC depends on sequence, interface chemistry, and three-dimensional geometry, but docking geometry is often summarized at the whole-receptor level, leaving CDR3-local pose difficult to compare across structures. We introduce TCR-FramePose, a local-frame descriptor set that represents each TCR–pMHC complex as three bodies — whole TCR, CDR3α, and CDR3β — measured relative to a pMHC groove frame. For each body, FramePose decomposes the native pose into reach, offset direction on S^2^, and orientation on *SO*(3); for tangent-space analyses, these components are mapped to six coordinates per body and 18 coordinates per complex.

Applied to 378 curated αβTCR–pMHC crystal structures, FramePose recovers known class-associated receptor-placement differences and additionally resolved whole-TCR and CDR3β orientation shifts that were not captured by crossing angle. The same orientation coordinates identified reverse-polarity and off-axis outliers as distinct modes. In cross-validated association analyses, FramePose added nonredundant BSA- and affinity-associated information beyond conventional descriptors, and the modest affinity gain was concentrated in CDR3 orientation blocks which were least recoverable from conventional descriptors.

Biological grouping analyses showed that shared receptor pose over peptide–MHC was organized primarily by germline V-region framework. TCRs recognizing the same peptide–MHC target favors shared FramePose geometries rather than strong receptor-specific divergence, whereas CDR3 sequence did not detectably reposition the rigid-body pose after antigen context and germline framework were fixed. MHC allele and peptide length contributed smaller adjustments, localized mainly to CDR3β and groove-normal orientation axes.

Finally, interface analyses showed that affinity tracked interface burial, with CDR3β reach linking FramePose geometry to binding through buried surface area. Within engineered panels, mutation-level effects were panel-specific, with CDR3β remodeling localizing to a recurrent interface region but varying in direction across receptors. These properties enable FramePose to serve as a geometric filter for in silico TCR–pMHC models and as a feature layer for structure-guided TCR engineering. Together, TCR-FramePose provides a nonredundant geometric layer for structure-guided TCR–pMHC analysis, linking germline-scaffolded recognition, CDR3-local pose, and interface organization without replacing sequence, contact, or energetic descriptors.

## Introduction

Recognition of peptide–major histocompatibility complex (pMHC) by αβ T cell receptors (TCRs) depends on molecular sequence, interface chemistry, and three-dimensional docking geometry^1–3^. Existing computational and structural approaches to TCR–pMHC recognition can be broadly grouped into three complementary strategies. The first is sequence-centered: motif analysis, biochemical-property features, repertoire similarity, and machine learning models seek to associate TCR, peptide, and MHC sequence with binding specificity^4–6^. These approaches are powerful for large-scale prediction, but they often do not specify how sequence variation is expressed structurally at the interface or how it changes receptor pose over the pMHC surface. The second is interface-energy-centered: structure-based modeling^7,8^, Rosetta scoring^9^, computational alanine scanning^2^, contact analysis^10–12^, and ΔΔ*G* estimation approaches^9,13^ to quantify how interfacial contacts and energetic terms contribute to binding. These approaches provide residue-level mechanistic information about the interface — for example, which residues contact one another, what types of interactions they form, and how mutations may alter binding energy-but they usually describe the interface after a pose has formed. The third is docking-pose-centered, geometric analyses^14–16^ describe where the TCR is placed over the pMHC surface and how it is oriented relative to the peptide-binding groove. This geometric layer is the most direct way to compare receptor placement and orientation across structures, but commonly used descriptors summarize the whole receptor and do not fully separate global TCR docking from local CDR3-loop pose.

We focus on docking pose because it is the structural layer that connects molecular determinants of binding to the physical organization of the TCR–pMHC interface^16–18^. A TCR variant may improve binding not only by adding favorable contacts, but also by reorienting the receptor so that existing contacts are brought into a more productive geometry^9,17,19,20^. Conversely, two complexes may have similar contact counts or similar global receptor positions but differ in how the receptor body or CDR3 loops are rotated over the pMHC platform^16,18,20,21^. Docking geometry is therefore not a secondary detail of recognition. Functional studies have shown that not every TCR–pMHC binding orientation is compatible with signaling^17,18^, and structural modeling frameworks increasingly treat TCR–pMHC docking as a rigid-body relationship between receptor and pMHC coordinate frames^8,14,16^. However, current structure-guided TCR analyses lack a standardized framework for describing docking pose as a decomposed geometric object. Existing descriptors can summarize receptor centering, projected docking angles, or whole-receptor frame transformations, but they do not provide a common coordinate system that separates receptor placement, receptor orientation, and local CDR3-loop placement and orientation.

TCR center-of-mass (TCR-CoM)^14,22^ descriptors describe the position of the TCR center relative to an MHC-centered coordinate system, typically through spherical coordinates. These descriptors are therefore primarily translational, in that they encode how far the receptor center lies from the pMHC reference and in what direction. The crossing angle^15,22^ and incident angle^22,23^ complement this by encoding receptor orientation, but they do so only as low-dimensional projected observables. Moreover, they are not pure orientation quantities. The crossing angle is the angle between the Vα–Vβ centroid line and the peptide axis after projecting the former onto the MHC groove plane, and the projection is itself sensitive to the in-plane position of the TCR centroid. The incident angle is defined as the tilt of the Vα–Vβ-to-peptide centroid-to-centroid vector relative to the groove normal and therefore changes whenever either centroid is translated even if the TCR’s orientation remains fixed. Crossing angle and incident angle together do not span the six rigid-body degrees of freedom and conflate orientation with translational variation by construction. Thus, an observed pose difference cannot be attributed unambiguously to groove-axis roll, cross-groove tilt, groove-normal twist, lateral or vertical receptor displacement, or a coupled change in receptor placement and orientation.

More recent rigid-body parameterizations, such as TCRdock^16^, provide a compact description of whole-TCR docking by representing the transformation between pMHC and TCR coordinate frames. TCRdock maps this transformation to a six-dimensional non-Euclidean descriptor set: the distance between frame origins, a torsion angle around the inter-origin axis, the unit direction from the pMHC frame to the TCR frame, and the reciprocal unit direction from the TCR frame back to the pMHC frame. This formulation is powerful for whole-receptor docking comparison and template-guided modeling. However, its parameters do not form a simple biological partition into translation and orientation. The distance and pMHC-frame unit direction describe receptor placement, whereas the reciprocal unit direction and torsion jointly encode aspects of receptor orientation relative to the line connecting the two frame origins. Thus, orientation information is distributed across reciprocal-direction and torsion coordinates rather than representing a full orientation block in fixed pMHC-aligned axes. In addition, these descriptors summarize the whole receptor and do not explicitly define local coordinate frames for CDR3α and CDR3β.

Here, we introduce TCR-FramePose, a local-frame descriptor set that represents TCR–pMHC geometry using three body frames — whole TCR, CDR3α, and CDR3β — measured relative to a common pMHC groove frame (Figure 1). For each body, FramePose defines the native pose by three components: reach, the Euclidean distance from the pMHC frame origin to the body-frame origin; offset, the unit direction of that displacement on S^2^; and orientation, the relative body-frame orientation on *SO*(3). Thus, the native FramePose representation preserves the natural geometry of translation and orientation. For statistical analyses requiring Euclidean coordinates, we map these native components to tangent space: one scalar reach coordinate, two S^2^ offset tangent coordinates, and three *SO*(3) orientation tangent coordinates per body, yielding an 18-coordinate tangent representation per complex. This design separates global receptor geometry from CDR3-local geometry while distinguishing how far each body sits from the groove, where it is displaced, and how it is oriented. A summary comparison of TCR-CoM, crossing/incident angle, TCRdock, and FramePose is provided in Supplementary Table S1.

**Figure 1.**
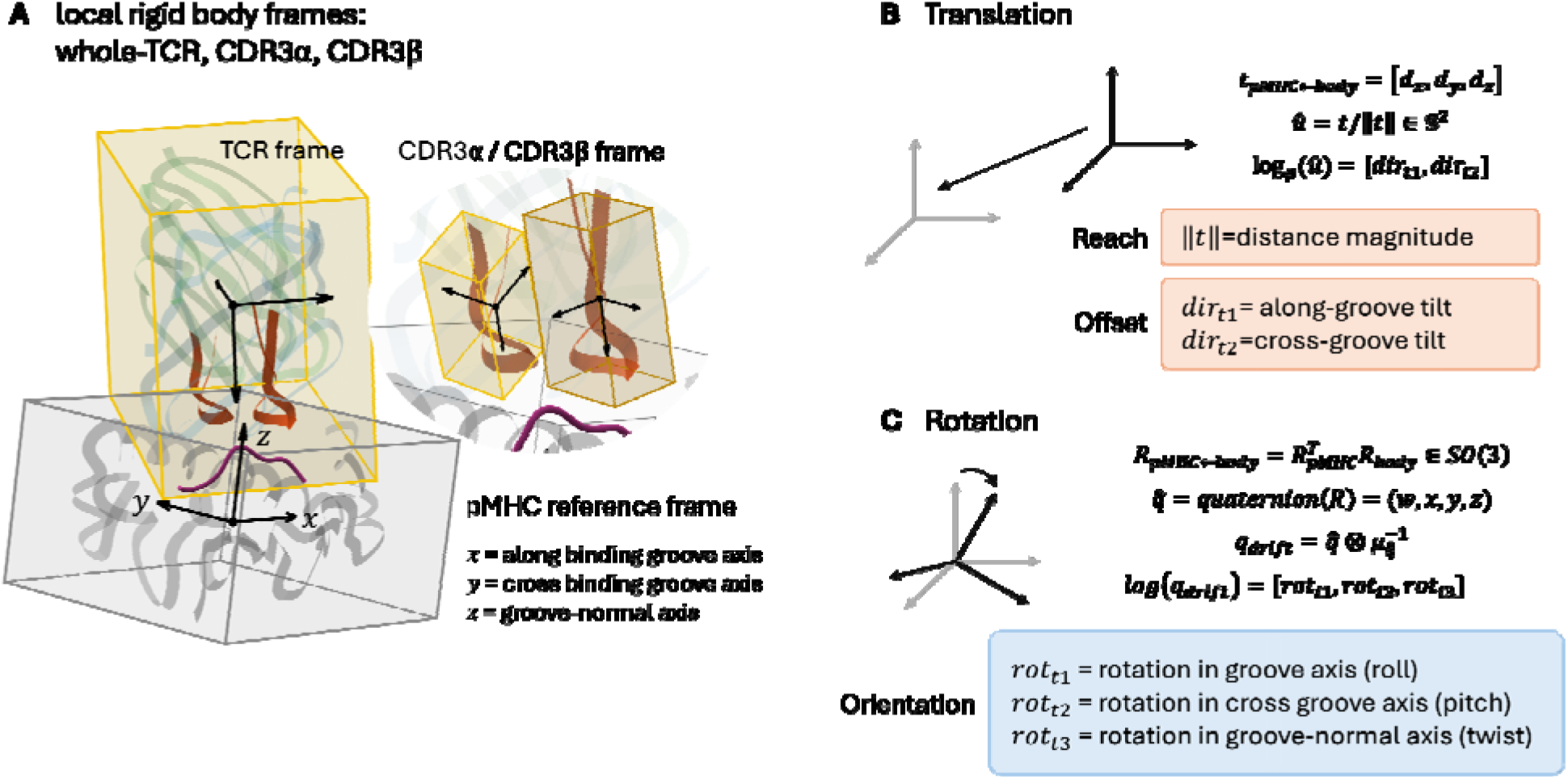
FramePose representation of global and CDR3-local TCR-pMHC docking geometry. FramePose represents each TCR–pMHC complex using three body frames — the whole TCR, CDR3α, and CDR3β — defined relative to a shared pMHC groove reference frame. (A) Local-frame construction. The pMHC reference frame is anchored to the peptide-binding groove, with the axes aligned along the groove (x), across the groove (y), and normal to the groove (z). Whole-TCR and CDR3 loop frames are defined relative to this reference. (B) Translational decomposition. The displacement from the pMHC origin to each body frame origin is decomposed into a reach (Euclidean distance, Å) and offset direction(unit vector). For statistical analysis, offset directions are log-mapped to two tangent coordinates ( and ) at the cohort Fréchet mean, corresponding approximately to along-groove and cross-groove tilt. (C) Orientation decomposition. Relative orientation between body and pMHC frame is represented as a rotation in and encoded as a unit quaternion. For statistical analysis, orientations are log-mapped to tangent coordinates ( , , ), corresponding to rotations about groove-aligned axes: groove-axis roll, cross-groove pitch, and groove-normal twist. Together, reach, offset, and orientation define decomposed global and local docking geometry.

Using curated αβTCR–pMHC crystal structures from TCR3d, we used FramePose to ask how global and local docking geometry is organized across structural, biological, and functional contexts. We first examined whether FramePose captures MHC class-associated pose variation and noncanonical polarity. We then tested whether FramePose contains BSA- or affinity-associated information beyond conventional docking descriptors. Moreover, we asked how biological labels and within-receptor perturbations shape FramePose space. Finally, we examined how FramePose geometry relates to interface burial and binding-associated structure, and whether engineered-panel remodeling reveals recurrent or receptor-specific CDR3β interface behavior. Together, these analyses evaluate whether separating reach, offset, and orientation across whole-TCR and CDR3 frames provides a useful geometric layer for structure-guided TCR–pMHC analysis.

## Methods

### Structural dataset

Human and mouse αβTCR–pMHC crystal structures were retrieved from the TCR3d database and filtered to a final cohort of 378 complexes (282 class I, 96 class II) with resolution ≤ 3.5 Å. Each retained structure carries the MHC peptide-binding platform, bound peptide, TCRα chain, and TCRβ chain. For class I complexes, the pMHC platform was defined by the MHC class I heavy chain; for class II complexes, both MHC class II α and β chains were required. TCRα/TCRβ chain identities and CDR annotations were assigned from the curated TCR3d metadata file and verified against the corresponding PDB chains. Only Cα atoms from standard amino-acid residues were used for geometric computations.

### FramePose representation and tangent coordinates

FramePose represents each TCR–pMHC complex as three local rigid body frames – the whole TCR variable domain and the CDR3α, and CDR3β loops - defined relative to a common pMHC groove reference frame. This design enables global receptor placement and CDR3-local geometry to be analyzed within a unified coordinate system (Figure 1).

For each body, pose relative to the pMHC frame is decomposed into translation and orientation. Translation vectors *t* were further decomposed into reach (distance) and offset direction. Reach was defined as *d*= ||*t*||, representing the Euclidean distance between the pMHC groove frame origin and the body-frame origin. Offset direction was defined as the unit vector *û* = *t*/||*t*||, where *û* ∈ S^2^. This decomposition separates how far a body lies from the groove (reach) from where it is positioned over the groove surface (offset direction). Orientation is defined as the relative rotation between the body frame and the pMHC frame, represented as a unit quaternion *q* ∈ *SO*(3).

For statistical analysis, offset directions and orientations were mapped to Euclidean tangent coordinates at cohort Fréchet means. Offset directions were log-mapped from S² to two tangent coordinates, *dir_t_*_1_ and *dir_t_*_2_, which describe directional deviation along and across the groove. Orientations were log-mapped from *SO*(3) to three tangent coordinates, *rot_t_*_1_, *rot_t_*_2_, and *rot_t_*_3_, representing rotations about groove-aligned axes. For interpretability, we refer to these components as groove-axis roll, cross-groove pitch, and groove-normal twist, respectively.

Each body contributes six tangent coordinates: one reach coordinate, two offset-direction coordinates, and three orientation coordinates. Across the three bodies — whole TCR, CDR3α, and CDR3β — this gives an 18-coordinate Euclidean tangent representation per complex.

Detailed frame construction procedures, including axis definitions, sign conventions, and numerical diagnostics are provided in Supplementary Note 2. Full derivations of the S^2^ and *SO*(3) log-maps, the Fréchet-mean estimation procedure, tangent-basis construction, and diagnostic assessments of tangent-plane validity are provided in Supplementary Note 3.

### Conventional global docking descriptor comparator

We compared FramePose against established global TCR–pMHC docking descriptors, including TCR-CoM^14^ spherical coordinates (r, θ, and ϕ); TCRdock^16^ parameters (distance *d*, torsion τ, and TCR/MHC unit-vector directions *û*_TCR➔ pM HC_ and *û*_pM HC➔ TCR_), crossing and incident angles^1^ (Supplementary Table S1). Consistent with the treatment of FramePose features, conventional descriptors defined on non-Euclidean spaces (θ, ϕ, τ, unit direction vectors, crossing angle, and incident angle) were mapped to tangent-space representation for multivariate analysis. Conventional-comparison analyses were restricted to complexes with complete FramePose and conventional descriptors (n = 377 for burial surface area and n = 244 for binding affinity association analyses).

To enable geometrically consistent comparison, conventional descriptors were grouped according to their underlying representation of rigid-body pose. TCR-CoM directional coordinates (θ, ϕ), defined as spherical coordinates of the TCR centroid relative to the pMHC origin, describe the direction of receptor placement over the groove and therefore correspond to the translational offset component of FramePose. In contrast, the crossing angle—defined as the angle between the projected Vα–Vβ axis and the peptide axis in the groove plane—captures a projected aspect of receptor orientation. Although crossing angle does not span the full rotational degrees of freedom in *SO*(3) and depends on centroid projection, it provides an observable related to receptor orientation and is therefore most directly comparable to FramePose orientation coordinates.

This mapping allows conventional descriptors to be compared to FramePose blocks in a manner that reflects their geometric meaning, rather than treating all descriptors as undifferentiated features.

### Descriptor distribution and class comparison

FramePose descriptor distributions were analyzed using native manifold geometry and tangent-space statistical tests.

Pairwise dependence between FramePose blocks was quantified using distance correlation computed from native distance matrices, with Euclidean distances for reach, and geodesic distances on S^2^ and *SO*(3) for offset and orientation, respectively. Statistical significance was assessed using 999 permutations with Benjamini–Hochberg correction.

Class-associated differences between MHC class I and class II complexes were quantified in two complementary ways. First, native class differences were reported as geodesic distances between class-specific Fréchet means, expressed in degrees for offset and orientation and in Å for reach. Second, statistical inference was performed in a shared tangent space at pooled Fréchet means.

In tangent space, scalar features (reach) were compared using Mann–Whitney U tests, and multicomponent features (offset and orientation) were compared using Hotelling’s *T*^2^ tests.

Effect sizes were reported as rank-based *r*^2^ or pseudo- = 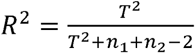, and p-values were corrected using the Benjamini–Hochberg procedure.

Orientation deviations for individual structures were interpreted using the norm of the *SO*(3) tangent vector, corresponding to geodesic rotation magnitude from the cohort mean, and further decomposed along groove-aligned axes.

### Cross-validated modeling framework

Associations between docking geometry and functional outcomes were evaluated using gradient-boosted decision tree models (HistGradientBoostingRegressor and Classifier). We used gradient boosting because tangent-coordinate relationships to biological outcomes may be nonlinear and interaction-dependent, especially because they summarize local embeddings of curved-manifold pose variables. Two association tasks were considered: buried surface area (BSA) regression (n = 378) and binding affinity (BA) classification using equilibrium dissociation constants *K_D_* (n = 244), where complexes were classified as strong binders (*K_D_* < 20 μM, n = 150) and weak binders (*K_D_* ≥ 20 μM, n = 94).

FramePose features were evaluated across a hierarchical model ladder consisting of single-body (whole TCR, CDR3α, CDR3β), two-body (TCR+CDR3α, TCR+CDR3β, CDR3α+CDR3β), and all-body configurations. For each task, the saturated FramePose model was defined as the configuration achieving the highest cross-validated performance. The all-body model was optimal for BSA regression, whereas the CDR3α+CDR3β model was optimal for affinity classification.

For comparison, conventional descriptor models were evaluated using TCR-CoM (5 features), crossing angle + incident angle (4 features), TCRdock (6 features), and composite 16 conventional descriptor set. This 16-feature model serves as the conventional baseline for augmentation analyses.

Models were evaluated using repeated five-fold cross-validation (20 repeats). Regression tasks used standard K-fold splits, and classification tasks used stratified K-fold splits. For each repeat, predictions were aggregated across folds, and performance metrics were computed from pooled out-of-fold (OOF) predictions. Regression performance was evaluated using coefficient of determination (*R*^2^) and Spearman correlation, and classification performance was evaluated using AUROC and AUPRC. All models used fixed hyperparameters (maximum iterations = 200, learning rate = 0.05, maximum tree depth = 4, minimum samples per leaf = 10, and L2 regularization = 1.0).

To localize contributions of geometric components, feature attribution analyses were performed on the task-specific saturated model. Leave-one-block-out analysis quantified model reliance by removing each FramePose block, while permutation importance quantified predictive signal by randomly permuting blocks in validation data.

To assess complementarity between descriptor families, augmentation analysis was performed by adding FramePose features to the conventional baseline model and measuring the resulting change in cross-validated performance.

Descriptor recoverability was evaluated by predicting FramePose features from conventional descriptors using repeated three-fold cross-validation (20 repeats), with performance measured using out-of-fold *R*^2^. The relationship between recoverability and augmentation gain was used to identify nonredundant geometric information captured by FramePose.

To further assess redundancy between descriptor families, we performed reverse-incremental analysis. In this framework, models were first trained using the saturated FramePose feature set for the task of interest. Conventional descriptor sets were then added on top of the saturated FramePose model. This analysis tests whether conventional descriptors provide additional association signal beyond that already captured by FramePose. Minimal or non-significant gains indicate that the corresponding conventional descriptors are largely redundant with the FramePose representation.

### Biological grouping and determinant analysis using partial PERMANOVA

To identify the biological determinants of TCR–pMHC docking geometry, we analyzed FramePose variation across the n = 378 αβTCR–pMHC cohort using a partial PERMANOVA framework applied to native FramePose distance matrices.

FramePose distances were computed at multiple levels. For each of the nine body-by-feature blocks (whole TCR, CDR3α, CDR3β × reach, offset, orientation), pairwise distances were calculated using the natural geometry of each component: Euclidean distance for reach, geodesic distance on S² for offset direction, and geodesic distance on *SO*(3) for orientation. Body-level distances were computed by averaging across the three feature components for each body, and a whole-pose composite distance was obtained by averaging across all nine blocks with mean normalization to ensure comparable contribution across components.

Biological grouping variables were derived from sequence-resolved annotations. Germline V-region identity was defined from framework and CDR1/2 regions of both α and β chains (excluding J-encoded FR4), while receptor identity combined germline V-region with CDR3 sequences. Additional variables included chain-specific V-region identity (α-V and β-V), MHC allele, peptide identity, peptide length, MHC class, and species.

Biological determinants were evaluated using Type-II partial PERMANOVA applied to distance matrices. Each determinant was tested using a (*focal* | *stratum*) design, in which the contribution of a focal factor (e.g., receptor identity, germline V-region, CDR3 sequence, MHC allele, or peptide features) was evaluated after accounting for variation explained by one or more conditioning variables (e.g., peptide–MHC or germline framework). The set of tested contrasts (e.g., receptor | peptide–MHC, V-region | allele, CDR3 | peptide–MHC + V-region, allele | V-region, peptide length | allele) is summarized in Table 2.

The partial variance attributable to a focal factor was computed using the difference in sum-of-squares as:

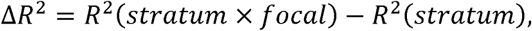

where *R*^2^ is derived from distance-based sums of squares. Adjusted Δ*R*^2^ values were reported to account for inflation associated with high-cardinality groupings.

Statistical significance was assessed using restricted permutation testing (N = 999), in which focal labels were permuted only within conditioning strata. This preserves the dependence structure induced by conditioning and ensures valid inference under confounded sampling. Permutation p-values were computed from the empirical null distribution, and Benjamini–Hochberg false discovery rate (FDR) correction was applied within each determinant family.

To contextualize effect sizes, an empirical reproducibility floor was estimated from true recrystallization pairs, defined as structures with identical TCR, peptide, and MHC sequences without engineered mutations or ambiguous residues. The median whole-pose composite distance among these structures (∼0.05) provided a baseline for minimal structural variation.

Because fine-grained groupings can inflate Δ*R*^2^ values through partitioning effects, results were interpreted relative to their restricted-permutation null distributions and compared across determinants with similar grouping granularity.

Determinant effects were evaluated at block level and coordinate (axis) level. At block level, we used native manifold distances for biologically interpretable components, whereas we used the 18 tangent coordinates to localize effects to specific geometric axes at axis level. Axis-level analyses enabled identification of directional contributions (e.g., groove-normal twist, cross-groove pitch) underlying each determinant.

### Interface burial, contact features, and cross-sectional FramePose–affinity models

To interpret how FramePose geometry relates to binding-associated interface structure, we analyzed affinity-annotated TCR–pMHC complexes using both cross-sectional and within-panel frameworks.

Affinity analyses were performed on a curated set of n = 244 structures with experimentally measured equilibrium dissociation constants (*K_D_*). Affinity values were compiled from TCR3d and ATLAS sources and curated to assign a single canonical *K_D_* per structure using standardized selection rules (Supplementary Note 7). Complexes lacking measurable *K_D_* values were excluded.

Structural interface properties were quantified using buried surface area (BSA), shape complementarity (SC), and contact-based features. Contact metrics were computed by aggregating interactions across CDR3α, CDR3β, and combined CDR3 loops, evaluated against both peptide-facing and MHC-facing regions, and stratified by interaction type, including hydrogen bonds, salt bridges, polar contacts, van der Waals packing, and total atomic contacts.

These combinations yielded 30 contact features, which were analyzed as raw counts in cross-sectional models.

FramePose predictors consisted of the 18 tangent coordinates representing reach, offset, and orientation for the whole TCR, CDR3α, and CDR3β bodies. Particular attention was given to CDR3β reach, defined as the Euclidean distance between the CDR3β frame origin and the pMHC groove frame. To distinguish within-allele variation from global structural differences, allele-referenced coordinates were computed by subtracting allele-specific means, and group-referenced tangent representations were used for directional variables when appropriate.

Cross-sectional relationships between structure and affinity were evaluated using linear mixed-effects models of the form:

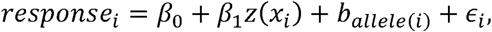

where the response was either BSA or log_10_ *K_D_, z*(*x_i_*) denotes the z-standardized predictor, and b*_allele_*_(*i*)_ is a random intercept for MHC allele to account for shared structural context. Separate models were fit for each interface descriptor and each FramePose coordinate, such that each coefficient *β*_1_ represents the change in response per standard deviation of the predictor. For affinity models, negative coefficients correspond to stronger binding. Model fitting was performed using maximum likelihood, and statistical significance was assessed using Wald tests with Benjamini–Hochberg false-discovery-rate correction applied within descriptor families.

To determine whether geometric effects reflect interface size rather than independent mechanisms, augmented models were fit including BSA and SC as covariates:

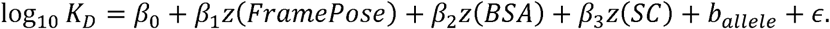

Comparison of coefficients across these nested models was used to assess whether pose–affinity associations persisted after accounting for interface burial and shape complementarity.

To evaluate the geometric linkage between pose and interface structure, a focused set of models was applied to CDR3β reach. Specifically, we tested (i) BSA as a function of reach, (ii) log_10_ *K_D_* as a function of reach, and (iii) log_10_ *K_D_* as a function of both reach and BSA within the same mixed-effects framework. These models assess whether reach is associated with affinity directly or indirectly through its relationship with interface burial. Because reach and BSA are geometrically coupled, these analyses were interpreted as structural associations rather than causal pathways.

To assess whether cross-sectional associations generalize to mutation-level effects, we analyzed engineered peptide panels in which receptor clonotype and MHC context are held constant while peptide sequence varies. Panels were defined by shared TCR identity and MHC allele and retained if at least two affinity-annotated structures were available, yielding 95 structures across 36 panels. Within each panel, affinity changes were represented relative to the strongest-binding structure using

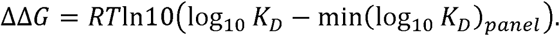

Structural changes were computed as panel-referenced differences. Contact remodeling was defined as change in contact features relative to panel means, whereas FramePose drift was defined as panel-referenced reach differences and tangent-space deviations for offset and orientation, computed using panel-level Fréchet means as base points.

Associations between structural changes and affinity differences were evaluated using mixed-effects models of the form

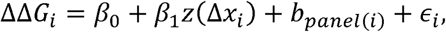

where Δ*x_i_* represents a panel-referenced structural feature and *b_panel_*_(*i*)_ is a panel-level random intercept. This formulation tests whether within-panel structural variation explains relative affinity differences while controlling for panel-specific baselines.

Statistical robustness of within-panel associations was assessed using both Benjamini–Hochberg correction and panel-restricted permutation testing, in which affinity values were permuted within panels while preserving panel membership. Features were considered robust only if they remained significant under both criteria.

For selected panels with sufficient sample size, residue-level CDR3β interface footprints were analyzed by computing minimum Cα–Cα distances between CDR3β residues and peptide or MHC contact-zone residues following structural superposition. Changes were evaluated relative to the strongest-binding structure within each panel. Because the number of suitable panels was limited, these analyses were interpreted as qualitative structural examples rather than pooled statistical tests.

## Results

### 1. FramePose resolves class-associated pose shifts and reverse-polarity outliers

We analyzed 378 crystallized αβ TCR–pMHC complexes from TCR3d (282 class I, 96 class II) using the FramePose representation (Figure 1). Each complex was represented relative to a common pMHC groove frame using three local body frames: the whole-TCR variable domain, CDR3α, and CDR3β.

For each body, pose was decomposed into translation and orientation. Translation was separated into reach, defined as the Euclidean distance from the pMHC frame origin to the body-frame origin, and offset direction, defined as the unit translation vector on S^2^. Orientation was defined as the relative rotation between the body and pMHC frames and represented as a unit quaternion. Thus, each body was described by three native geometric components: reach, offset direction, and orientation.

For statistical analysis, manifold-valued components were mapped to tangent space at cohort Fréchet means. Offset directions were projected to two tangent coordinates describing displacement along and across the groove, and orientations were projected to three coordinates corresponding to groove-axis roll, cross-groove pitch, and groove-normal twist. Each body therefore contributed six coordinates, yielding an 18-dimensional representation per complex.

All constructed frames satisfied geometric consistency criteria, including orthogonality, right-handedness, and stability diagnostics (Supplementary Note 2; Supplementary Figure S1).

We first examined the internal structure of the FramePose representation. Pairwise distance correlations, computed using native manifold distances (Euclidean for reach, geodesic for offset and orientation), showed strong cross-body coupling of offset-direction coordinates across whole-TCR, CDR3α, and CDR3β frames (Supplementary Figure S2). This coupling reflects the physical linkage of CDR3 loops to the shared TCR variable-domain body.

We next asked whether FramePose captures class-associated differences in docking geometry. Previous studies have shown that MHC class differences are reflected in TCR placement over the pMHC surface, particularly in TCR-CoM directional coordinates, whereas crossing angle alone often fails to distinguish class I and class II docking modes (Figure 2A).

**Figure 2.**
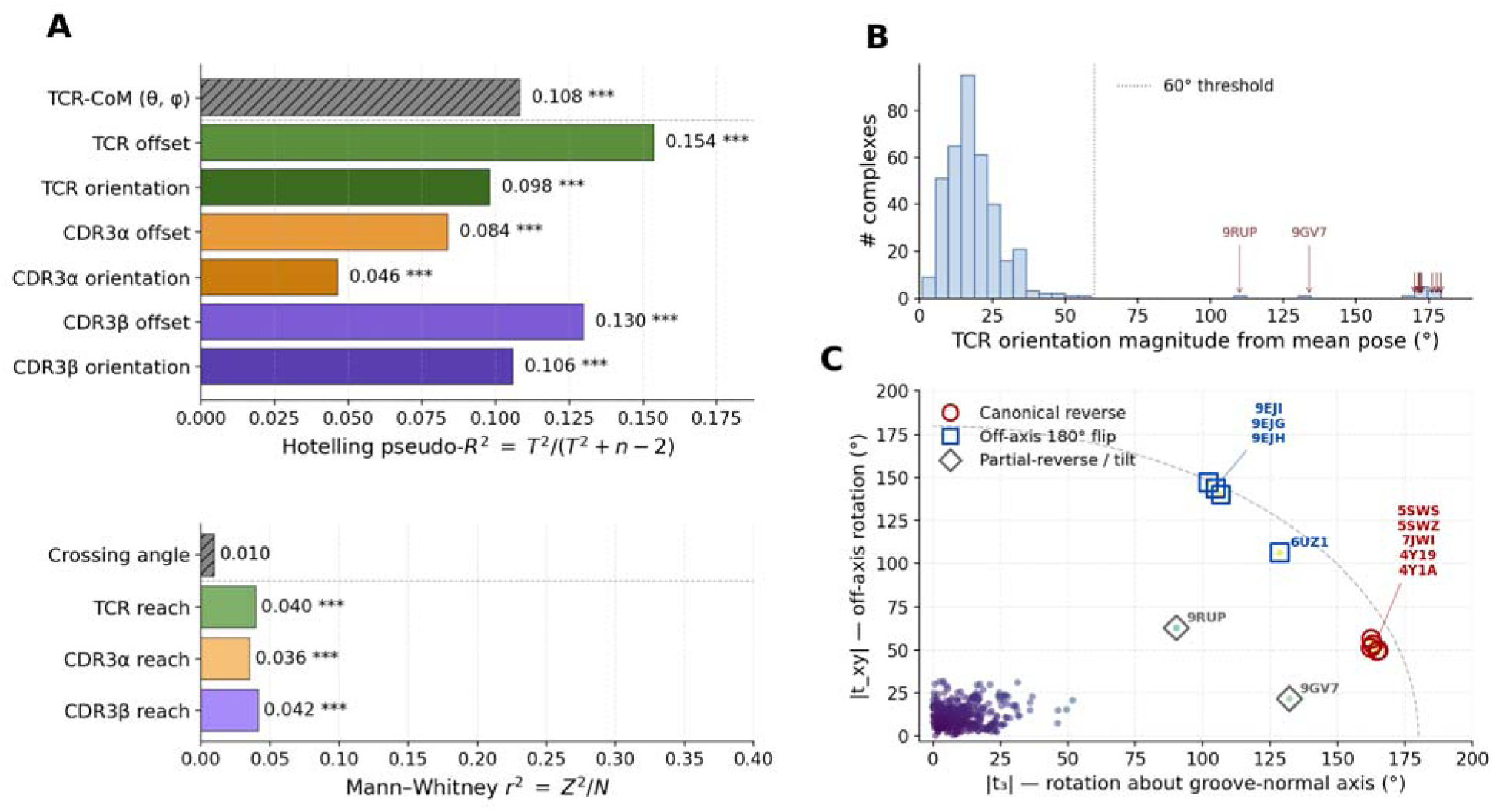
FramePose resolves class-associated docking differences and orientation outliers. FramePose descriptors were computed for n = 378 αβTCR–pMHC structures (282 class I, 96 class II). (A) Class-associated differences. Separation between class I and class II complexes is shown for conventional docking descriptors and FramePose blocks. Multicomponent features (offset on and orientation on , as well as TCR-CoM directional coordinates) were tested using Hotelling’s tests in tangent space at the pooled Fréchet mean, with effect size reported as pseudo- = . Scalar features (reach and crossing angle) were tested using Mann–Whitney U tests with rank-based effect size . -values are Benjamini–Hochberg FDR-corrected (*** q < 0.001; ** q < 0.01; n.s., not significant). FramePose recovers known class-associated translational differences and additionally resolves orientation differences not captured by crossing angle. (B) Whole-TCR orientation deviation. Distribution of SO(3) geodesic rotation distances (degrees) between each complex’s whole-TCR orientation and the cohort Fréchet-mean orientation. Most complexes lie within ∼60°, corresponding to canonical docking, whereas a subset shows large deviations (>110°), indicating noncanonical orientations. (C) Outlier decomposition. Orientation outliers are decomposed in tangent space into groove-normal twist ( ) and off-axis rotation ( ). The dashed curve indicates total rotation of 180°. These components distinguish canonical reverse-polarity docking, near-180° off-axis rotations, and partial-reverse or tilted geometries.

Because TCR-CoM directional coordinates (θ, ϕ) describe receptor placement over the pMHC surface, they serve as a natural comparator for the translational component of FramePose. In contrast, crossing angle captures a projected aspect of receptor orientation—defined relative to the groove plane—and therefore provides an orientation-related but incomplete observable.

Consistent with prior work, TCR-CoM directional coordinates (θ, ϕ) showed significant class separation (Hotelling’s *T*^2^, pseudo *R*^2^=0.108, p<0.00), whereas crossing angle showed minimal separation (Figure 2A). This difference is consistent with underlying geometry: class I and class II MHC molecules differ in binding groove architecture^1,24^, leading to systematic differences in receptor placement, while crossing angle captures only a single projected dimension of orientation and does not span the full rotational degrees of freedom.

FramePose recapitulated this translational shift and revealed additional orientation differences. Native manifold comparisons showed mean class-associated offset-direction tilts of approximately 10° and orientation differences of approximately 12–18° across body frames (Table 1). Statistical testing was performed in tangent space to localize these effects, using

**Table 1.**
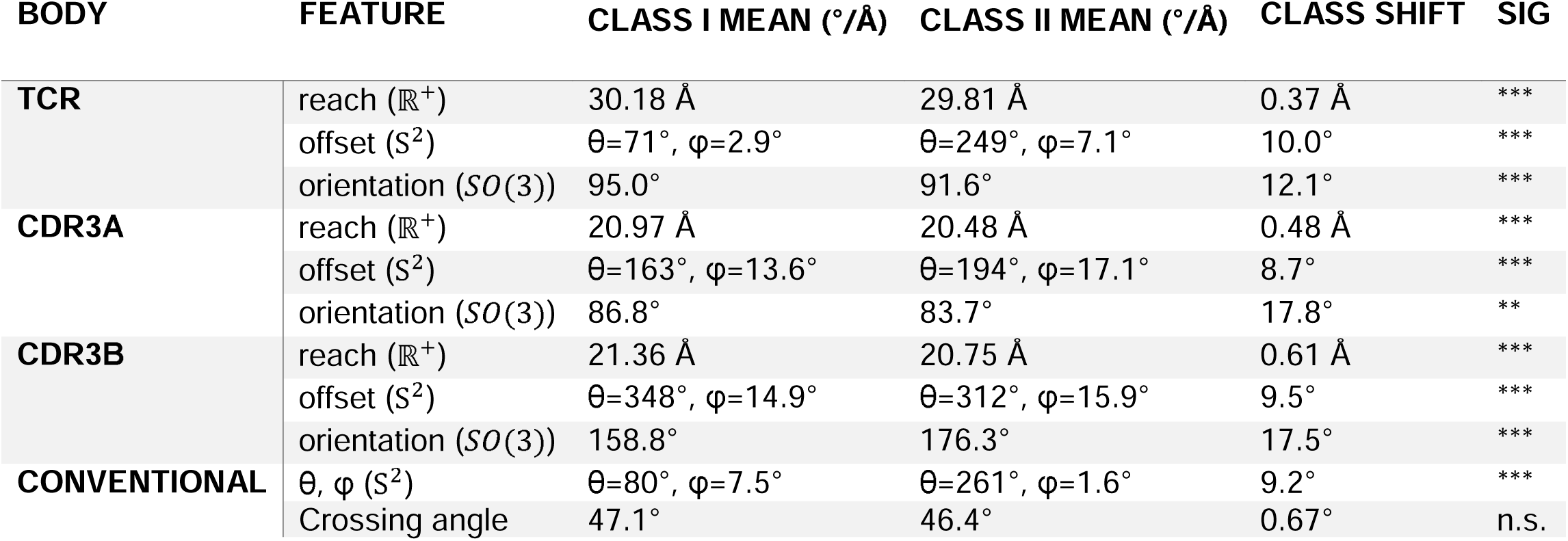
Class-associated shifts in TCR–pMHC docking geometry expressed in native geometric units. Mean poses of the whole TCR, CDR3α, and CDR3β frames are compared between MHC class I (n = 282) and class II (n = 96) complexes. Each feature is evaluated in its native geometric space: reach as a Euclidean distance (Å), offset as a unit direction on S², and orientation as a rotation on *SO*(3). Class-specific means are computed as follows: arithmetic mean for reach, and Fréchet (Karcher) means for offset directions and orientations. Offset means are reported as spherical angles (θ, ϕ; degrees), and orientation means are summarized by geodesic rotation angles (degrees). Class shift is defined as the geodesic distance between class-specific means on the corresponding manifold (Å for reach; degrees for offset and orientation). Statistical significance is assessed in a shared tangent space at the pooled Fréchet mean: scalar features (reach, crossing angle) using two-sided Mann–Whitney U tests, and multicomponent features (offset, orientation, TCR-CoM) using Hotelling’s *T*² tests. Effect sizes for multicomponent tests are reported as pseudo-*R*² = *T*² / (*T*² + *n*₁ + *n*₂ - 2). P-values are Benjamini–Hochberg false discovery rate–corrected (*** q < 0.001; ** q < 0.01; n.s., not significant).

| BODY | FEATURE | CLASS I MEAN ($^\circ/\text{\AA}$) | CLASS II MEAN ($^\circ/\text{\AA}$) | CLASS SHIFT | SIG |
| --- | --- | --- | --- | --- | --- |
| TCR | reach ($\mathbb{R}^+$) | 30.18 $\text{\AA}$ | 29.81 $\text{\AA}$ | 0.37 $\text{\AA}$ | *** |
| | offset ($S^2$) | $\theta=71^\circ, \phi=2.9^\circ$ | $\theta=249^\circ, \phi=7.1^\circ$ | 10.0$^\circ$ | *** |
| | orientation ($SO(3)$) | 95.0$^\circ$ | 91.6$^\circ$ | 12.1$^\circ$ | *** |
| CDR3A | reach ($\mathbb{R}^+$) | 20.97 $\text{\AA}$ | 20.48 $\text{\AA}$ | 0.48 $\text{\AA}$ | *** |
| | offset ($S^2$) | $\theta=163^\circ, \phi=13.6^\circ$ | $\theta=194^\circ, \phi=17.1^\circ$ | 8.7$^\circ$ | *** |
| | orientation ($SO(3)$) | 86.8$^\circ$ | 83.7$^\circ$ | 17.8$^\circ$ | ** |
| CDR3B | reach ($\mathbb{R}^+$) | 21.36 $\text{\AA}$ | 20.75 $\text{\AA}$ | 0.61 $\text{\AA}$ | *** |
| | offset ($S^2$) | $\theta=348^\circ, \phi=14.9^\circ$ | $\theta=312^\circ, \phi=15.9^\circ$ | 9.5$^\circ$ | *** |
| | orientation ($SO(3)$) | 158.8$^\circ$ | 176.3$^\circ$ | 17.5$^\circ$ | *** |
| CONVENTIONAL | $\theta, \phi$ ($S^2$) | $\theta=80^\circ, \phi=7.5^\circ$ | $\theta=261^\circ, \phi=1.6^\circ$ | 9.2$^\circ$ | *** |
| | Crossing angle | 47.1$^\circ$ | 46.4$^\circ$ | 0.67$^\circ$ | n.s. |

Mann–Whitney tests for scalar reach and Hotelling’s *T*^2^ tests for multicomponent offset and orientation blocks. The strongest class discriminator was whole-TCR offset direction, with a 9.99° mean tilt and pseudo-*R*² = 0.154 (Table 1, Figure 2A) exceeding TCR-CoM (θ, ϕ) separation. CDR3β offset direction also showed substantial class-associated displacement, with a 9.48° mean tilt and pseudo-*R*² = 0.129 (Table 1, Figure 2A). These results indicate that class-associated receptor repositioning is reflected both globally and at the CDRβ loop level.

Importantly, orientation differences ranked among the next strongest signals. CDR3β and whole-TCR orientation showed mean rotations of 17.54° and 12.15°, with pseudo-*R*² = 0.104 and 0.094, respectively (Table 1, Figure 2A). These effects were not captured by crossing angle, which exhibited minimal class separation (47.1° vs 46.4°, r²=0.01, p=n.s.; Table 1, Figure 2A). FramePose therefore reproduces the class-associated receptor-placement difference captured by TCR-CoM while additionally resolving class-associated whole-TCR and CDR3β orientation differences that are not captured by crossing angle alone.

The orientation tangent coordinates also identified noncanonical docking modes at the level of individual complexes. The norm of the whole-TCR orientation tangent vector quantifies the geodesic rotation distance from the cohort Fréchet-mean orientation. Most complexes exhibited rotation magnitudes below 60°, whereas a small subset showed large deviations between ∼110° and ∼180° (Figure 2B).

Decomposition of the orientation vectors into groove-normal (twist) and off-axis components distinguished different outlier modes (Figure 2C). Five complexes formed a canonical reverse-polarity cluster: 5SWS, 5SWZ^25^, 7JWI^18^ representing TRBV17⁺ NP-specific TCRs on H2-Db, and 4Y19 and 4Y1A^26^ representing FS17/FS18 on HLA-DR4/proinsulin. These complexes showed ∼170° rotation dominated by groove-normal twist (94–96% of rotation magnitude; Figure 2B-C), consistent to the canonical definition of reverse-polarity docking. Despite spanning different MHC classes, these five structures collapsed into a tight single cluster in the FramePose whole-TCR orientation space, indicating that they share the same kinematic polarity reversal despite belonging to different MHC classes (Figure 2C).

This clustering was less apparent in the conventional descriptor projection. In crossing angle vs TCR-CoM ϕ space, the same five complexes separated into class-specific subclusters: the three H2-Db class I complexes near ϕ ∼ 50° and the two HLA-DR4 class II complexes near ϕ ∼ 12° (Supplementary Figure S4B). This separation likely reflects class-dependent differences in the platform geometry that affect centroid-based positioning, rather than a distinct polarity mechanism. By representing relative orientation as a *SO*(3) tangent vector, FramePose groups the shared reverse-polarity mode independently of these centroid-position differences.

A second cluster of four complexes - 9EJI, 9EJG, and 9EJH^27^ representing peptide-agnostic G9 TCR on HLA-DQ2.5 together with 6UZ1^28^, an engineered A6 affinity variant - also showed TCR rotated ∼ 180° from the cohort Fréchet-mean pose (Figure 2B). However, only 57-77% of each rotation vector was aligned with the groove-normal axis, with the remaining component distributed across the off-axis plane (Figure 2C). These structures represent off-axis 180° flip rather than canonical groove-normal polarity flips.

Three additional complexes, 6D7G, 9GV7, and 9RUP, showed intermediate rotation magnitude (110°–152°; Figure 2B) and majority rotated along the groove-normal axes (0.82–0.99; Figure 2C), corresponding to partial-reverse or tilted docking mode. Across all outlier groups, CDR3α and CDR3β orientation deviation tracked whole-TCR rotation norm (Supplementary Table S2), suggesting that outlier signal reflects a whole-receptor polarity change rather than isolated loop distortion.

Together, these analyses show that FramePose captures both known class-associated docking differences and previously unresolved orientation variation. The same orientation coordinates used for class-level analysis also provide a quantitative framework for identifying and classifying noncanonical docking geometries.

### 2. FramePose adds CDR3-local information for interface size, but rigid-body pose alone only weakly captures binding affinity

We next asked whether FramePose geometry was associated with structural and functional readouts, and whether FramePose-associated signal overlapped with conventional docking descriptors. We considered two outcomes — buried surface area (BSA), a continuous structural readout of interface size, and a binary binding-affinity label derived from *K_D_*.

FramePose was evaluated across hierarchical body configurations, including single-body, two-body, and all-body models. For BSA, the saturated FramePose model was the all-body configuration, whereas for affinity classification, the saturated model consisted of the CDR3α and CDR3β bodies. Conventional descriptors included TCR-CoM, TCRdock, crossing/incident angles, and their combined 16-feature baseline (Supplementary Table S1).

#### 2.1 CDR3-local orientation accounts for the FramePose-specific association with BSA

We first evaluated association with buried surface area (BSA; n = 377 after requiring complete conventional descriptors). Across the FramePose model hierarchy, the all-body configuration showed the strong association (*R*^2^= 0.5; Figure 3A), identifying it as the saturated model for BSA regression. Single-loop models were weaker (CDR3α 0.348, CDR3β 0.379), although combining CDR3α and CDR3β substantially improved association strength (0.48), indicating complementary loop-level contributions.

**Figure 3.**
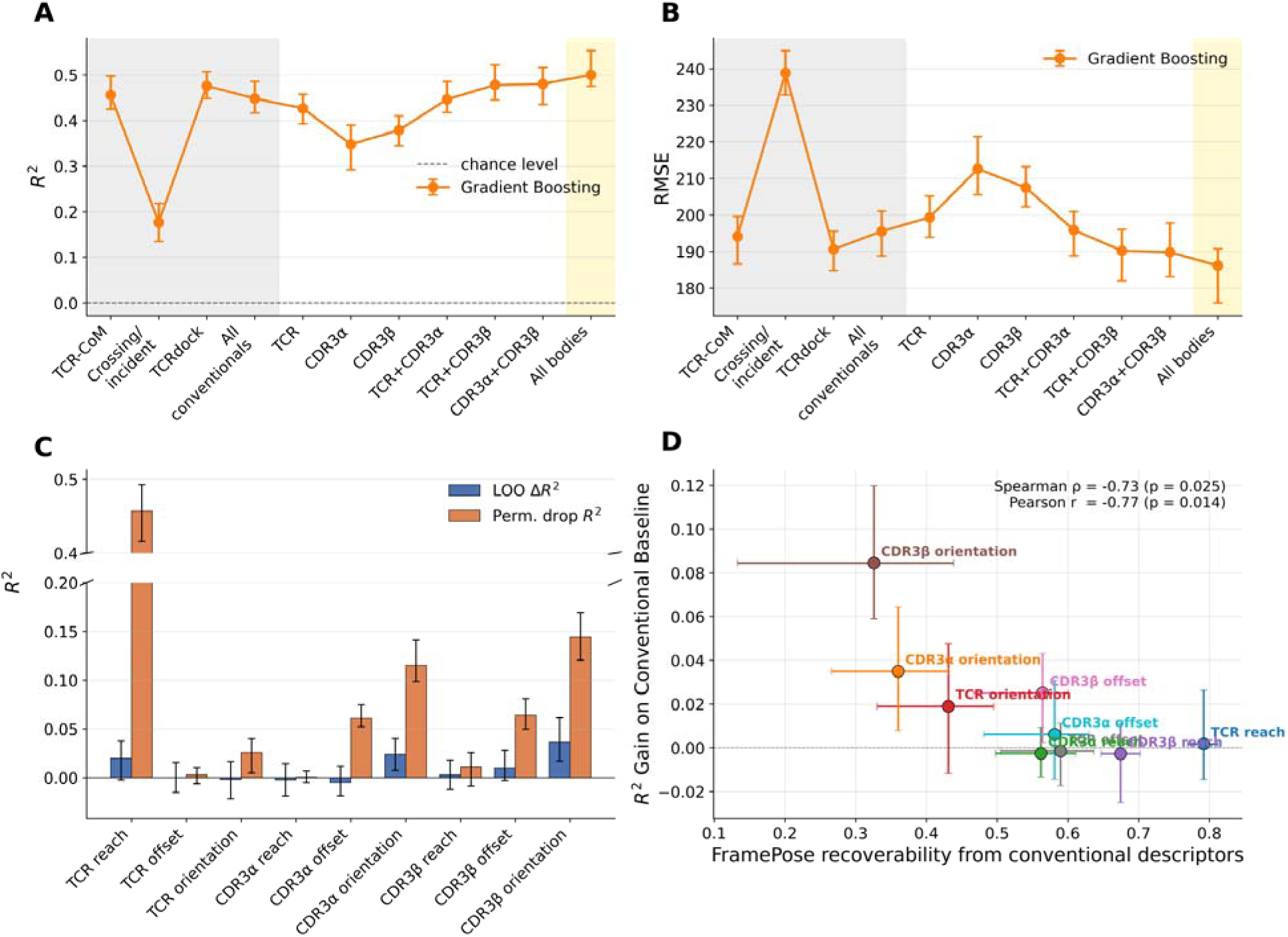
FramePose contributions to burial surface area modeling. Analyses were performed on n = 377 complexes with complete conventional docking descriptors. (A–B) Association performance. Cross-validated association between docking geometry and buried surface area (BSA, Å²) was quantified using gradient boosting regression models and evaluated using coefficient of determination ( ) and root mean squared error (RMSE). Values are reported as the mean and empirical interval across 20 repeated cross-validation splits. Models included conventional families (grey blocks), and the hierarchical FramePose model ladder. The all-bodies FramePose model achieves the highest association and is used as the saturated model (yellow block). (C) Block contributions. Within the saturated all-bodies gradient-boosting model, block importance is assessed by leave-one-block-out analysis (LOO ), defined as the reduction in association strength when a block is removed from training, and permutation importance (Perm. drop ), defined as the reduction after random shuffling of block features in held-out folds while preserving marginal distributions. (D) Augmentation and nonredundancy. Each FramePose block is characterized by its recoverability from conventional descriptors (cross-validated for predicting the block from conventional features) and its augmentation gain ( ) when added to a conventional baseline model. Spearman and Pearson correlations quantify the relationship between these quantities. Blocks with low recoverability—particularly CDR3 orientation features—show greater augmentation gains, indicating that they provide association signal beyond conventional descriptor representations.

Conventional descriptors captured much of the global BSA-associated signal. TCRdock was the strongest individual family (R² = 0.476), and conventional baseline achieved comparable to TCR-only FramePose model (∼0.43∼0.45; Figure 3A), indicating that much of the global BSA-associated signal is captured by whole-receptor placement alone.

Within the saturated FramePose model, feature attribution showed that whole-TCR reach was the largest contributor (permutation drop Δ*R*^2^ = 0.457; Figure 3C). However, CDR3-local components also contributed consistently, with orientation of both loops forming the next strongest tier (permutation drop 0.115 and 0.144 for CDR3α and CDR3β, respectively) followed by small contributions from offset features(permutation drops 0.061 and 0.064 for CDR3α and CDR3β, respectively). Thus, BSA-associated information is distributed across both global TCR placement and CDR3-local orientation, rather than concentrated in a single cluster of correlated coordinates.

To isolate FramePose-specific contribution, we performed augmentation analysis relative to the conventional baseline (Figure 3D). A subset of CDR3-local features increased association strength, most notably CDR3β orientation (Δ*R*² = 0.085), CDR3α orientation (Δ*R*² = 0.035), and CDR3β offset (ΔR² = 0.025). In contrast, whole-TCR reach and offset provided little additional gain, consistent with their global-placement signal already largely captured by conventional centroid-based descriptors. These results indicate that while global receptor placement accounts for much of the overall BSA-associated signal, CDR3 orientation provides additional association signal beyond conventional docking representations.

#### 2.2 Affinity-associated FramePose signal is modest and concentrated in CDR3-local geometry

We next evaluated association with binding-affinity class (n = 244; 150 strong, 94 weak). Binding-affinity classification showed substantially weaker association strength than BSA regression. Across the conventional descriptor sets, AUROC remained near chance (0.53–0.54), and AUPRC (0.64–0.66) only marginally exceeded the class baseline (Figure 4A).

**Figure 4.**
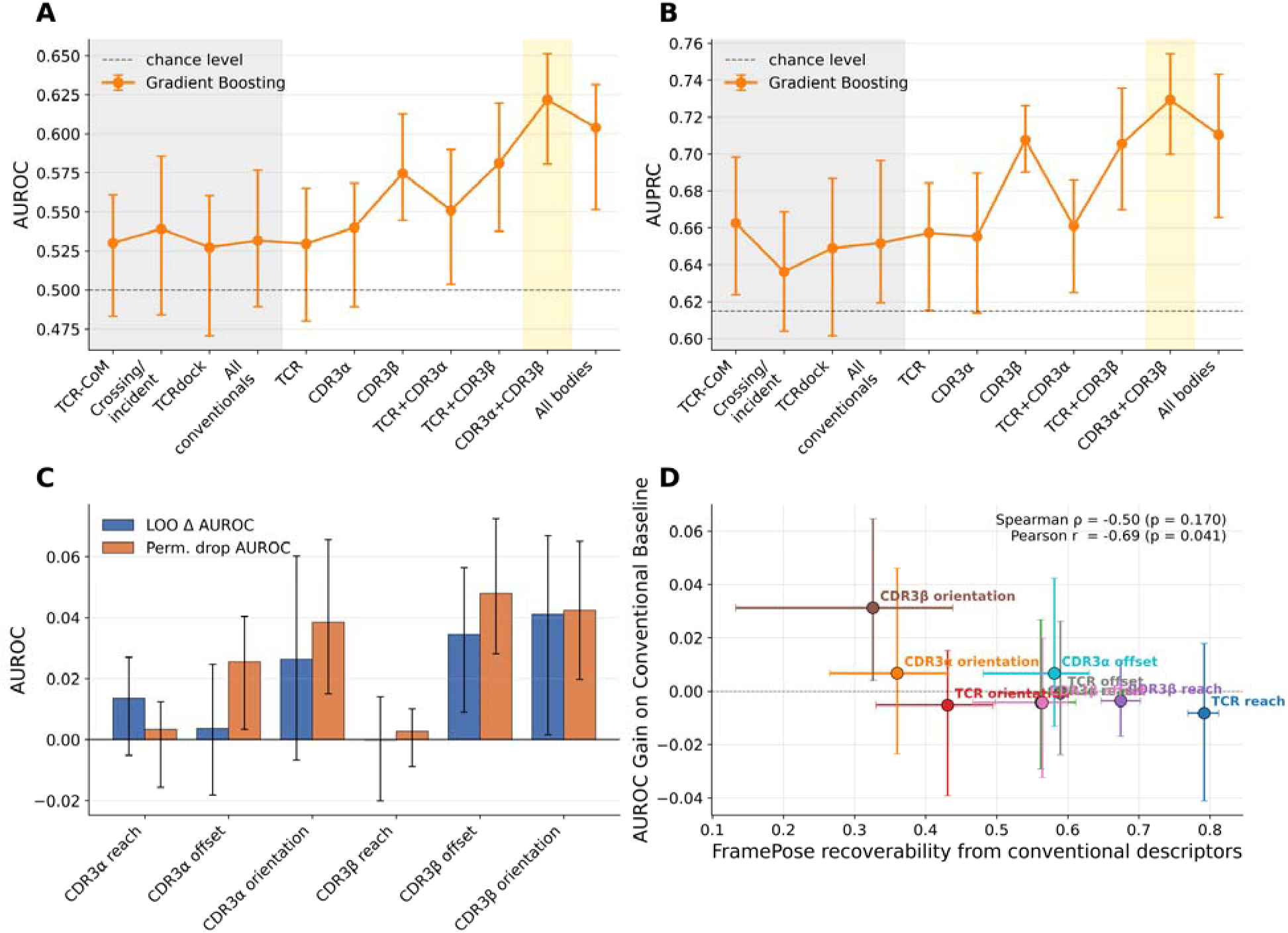
CDR3-local FramePose features show modest association with binary binding-affinity labels. Analyses were performed on n = 244 complexes with curated dissociation constants ( , μM). Complexes were classified as strong binders ( < 20 μM, n=150) or weak binders ( ≥ 20 μM, n=94). (A–B) Association performance. Cross-validated association between docking geometry and binding affinity ( ) was quantified using gradient boosting classification models and evaluated using AUROC and AUPRC. Values are reported as the mean and empirical interval across 20 repeated cross-validation splits. Chance levels are 0.5 (AUROC) and the positive-class fraction (AUPRC ≈ 0.615). Models included conventional families (grey blocks), and the hierarchical FramePose model ladder. The CDR3α+CDR3β FramePose model achieves the highest association and is used as the saturated model (yellow block). (C) Block contributions. Within the saturated all-bodies gradient-boosting model, block importance is assessed by leave-one-block-out analysis (LOO AUROC), defined as the reduction in association strength when a block is removed from training, and permutation importance (Perm. drop AUROC), defined as the reduction after random shuffling of block features in held-out folds while preserving marginal distributions. CDR3-local orientation and offset blocks contribute more strongly than reach. (D) Augmentation and nonredundancy. Each FramePose block is characterized by its recoverability from conventional descriptors (cross-validated for predicting the block from conventional features) and its augmentation gain ( AUROC) when added to a conventional baseline model. Spearman and Pearson correlations quantify the relationship between these quantities. Blocks with low recoverability—particularly CDR3 orientation features—show greater augmentation gains, indicating that they provide association signal beyond conventional descriptor representations.

FramePose showed stronger association with affinity labels within this cohort, although overall association strength remained modest. The saturated CDR3α+CDR3β model reached AUROC = 0.622 and AUPRC = 0.729, exceeding conventional baseline with a 0.09 AUROC gain and 0.08 AUPRC gain (Figure 4A,B). Association signal was concentrated in CDR3-local geometry, that CDR3-only model outperformed the all-body configuration and inclusion of whole-TCR features did not improve association beyond CDR3 loops, indicating limited contribution from global receptor geometry once local features were included. The two CDR3 loops carried complementary information: CDR3α alone reached AUROC = 0.54, CDR3β alone reached AUROC = 0.575, and combining them increased AUROC to 0.622.

Within the saturated CDR3-only FramePose model, orientation was the geometric component most consistently associated with affinity, across both CDR3 loops and both sensitivity analyses. CDR3β orientation was the single dominant block, that its removal produced a largest reduction in association strength (LOO ΔAUROC = 0.041) and it remained a top contributor under permutation (permutation ΔAUROC = 0.043; Figure 4C). CDR3α orientation was the next most consistent block (LOO ΔAUROC = 0.026; permutation ΔAUROC = 0.039), so that the two rotational blocks together accounted for the majority of the recoverable orientation–offset signal. Offset contributed secondarily and asymmetrically: it was carried almost entirely by CDR3β offset (permutation ΔAUROC = 0.048; LOO ΔAUROC = 0.035), whereas CDR3α offset was weak and reached significance only under permutation (permutation ΔAUROC = 0.026). The CDR3 reach blocks of both loops were uninformative across all sensitivity analyses. The affinity-associated pose signal is therefore carried predominantly by how the CDR3 loops are rotationally oriented, rather than by how far they reach.

These results indicate that rigid-body pose alone is not sufficient to explain binding affinity. Even the strongest FramePose models remained in a modest AUROC range, and the *K_D_* values were heterogeneous and binarized. Nevertheless, the analysis identifies a reproducible but limited CDR3-local pose signal, most consistently involving CDR3β orientation, with secondary contributions from CDR3β offset and CDR3α orientation. This signal was not captured by conventional global docking summaries, but accurate affinity modeling will likely require sequence-level, energetic, contact-level, or atomistic features beyond rigid-body FramePose descriptors alone.

### 2.3 Limited recoverability of CDR3 orientation explains its strong association contribution

To understand why the BSA- and affinity-associated FramePose signals concentrated in CDR3-local coordinates, we evaluated recoverability of FramePose features from conventional descriptors (Supplementary Figure S6). Recoverability was quantified as out-of-fold *R*^2^ for multicomponent targets, providing a measure of how well each FramePose block can be reconstructed from conventional docking summaries.

As expected, global translational features were highly recoverable. Whole-TCR reach showed highest recoverability (*R*^2^=0.79; Supplementary Figure S6A), indicating that conventional descriptors jointly capture most variation in receptor distance to the pMHC frame. Offset and reach for CDR3 showed intermediate recoverability along with TCR offset (0.56∼0.67; Supplementary Figure S6A).

In contrast, orientation blocks were the least recoverable features. Across FramePose blocks, augmentation gain decreased with increasing recoverability, such that poorly recovered features contributed most strongly to association models (Figure 3D, Figure 4D). The least recoverable block CDR3β orientation provided largest gains in both BSA and affinity association analyses (BSA Δ*R*^2^ = 0.084 [0.059, 0.120]; affinity ΔAUROC = 0.031 [0.004, 0.065]), whereas highly recoverable features such as whole-TCR reach (recoverability *R*^2^= 0.792) showed near-zero marginal contribution (BSA Δ*R*^2^ = 0.002 [−0.014, 0.026]; affinity ΔAUROC = −0.008 [−0.041, 0.018]). Reverse augmentation analysis yielded consistent results, with minimal gains when conventional descriptors were added to saturated FramePose models (Supplementary Figure S5).

This pattern explains why FramePose-specific BSA and affinity association signals were localized to CDR3 orientation blocks. Features that are poorly captured by conventional descriptors, particularly CDR3-local orientation, contribute more strongly than would be expected from their recoverability, identifying them as the primary source of nonredundant geometric information in FramePose.

### 3. Germline V-region framework organizes shared receptor pose over peptide–MHC

We next asked how biological identity organizes docking geometry. Specifically, we tested whether TCRs recognizing the same peptide–MHC (pMHC) adopt receptor-specific poses or instead converging toward shared docking geometries, and which sequence or antigen features determine this organization. This distinction is important because multiple distinct TCR clonotypes can recognize the same peptide–MHC target while retaining substantial CDR3 diversity^29–32^, yet their overall receptor placement and orientation may remain constrained by germline V-region framework^17,24,33,34^.

Across our structural cohort, receptor, peptide, and MHC labels are strongly confounded, that a receptor is usually crystallized on a single allele with one or a few peptides, and shared antigens often recruit biased V-gene usage. We therefore used a sequence-resolved, conditioned PERMANOVA design to separate the contributions of germline V-region framework, CDR3 sequence, MHC allele, and peptide features to FramePose variation (Supplementary Table S3). This approach tests each determinant within matched strata, enabling separation of germline, antigen, and junctional-loop effects on FramePose geometry.

#### 3.1 TCRs recognizing the same peptide–MHC share germline-anchored receptor geometry

We first tested whether different receptors recognizing the same pMHC retain distinct docking poses. Within matched pMHC contexts, receptor identity explained little residual whole-pose variation and was not significant (receptor | pMHC, adjusted Δ*R*^2^= 0.087, p = 0.136; Figure 5B; Table 2). A non-significant effect cannot by itself prove convergence, but it shows that TCRs recognizing the same pMHC target did not accompanied strong receptor-specific pose divergence in the available cohort.

**Figure 5.**
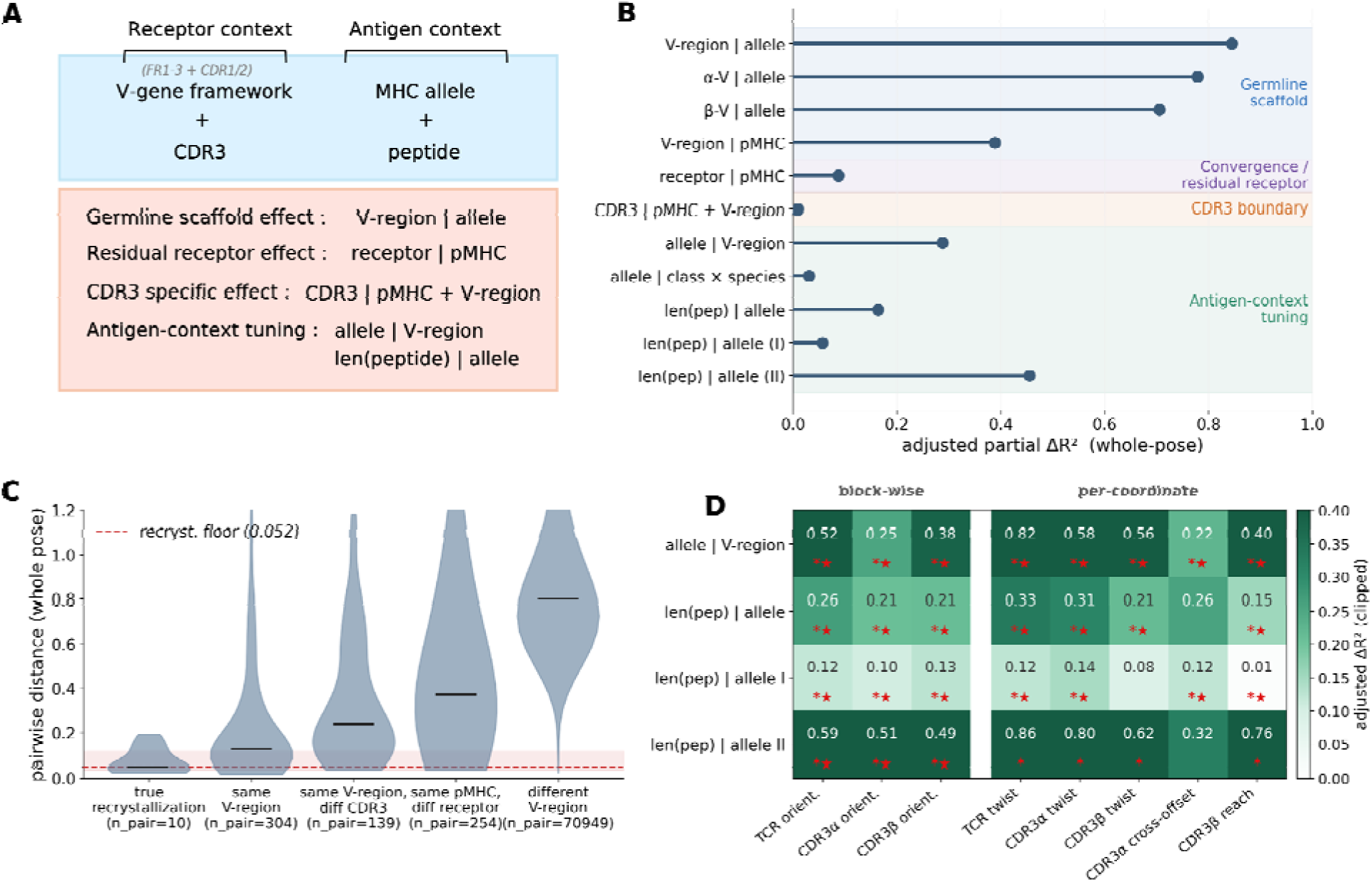
Germline V-region framework organizes shared TCR–pMHC pose geometry. Docking-geometry was analyzed across n=378 structures using sequence-derived grouping variables and conditioned partial PERMANOVA applied to native FramePose distance matrices. (A) Conditioned contrast design. Each analysis is formulated as a conditioned contrast of the form , in which the effect of a biological determinant (focal) is evaluated after accounting for variation explained by matched strata. Focal labels are permuted only within conditioning groups to preserve the structure of the comparison. (B) Whole-pose determinant hierarchy. Adjusted values from partial PERMANOVA quantify the fraction of variance in whole-pose geometry attributable to each determinant beyond conditioning variables. Germline V-region identity shows the strongest association with docking geometry (V-region | allele), whereas receptor identity within fixed peptide–MHC context (receptor | pMHC) and CDR3 sequence conditioned on antigen and germline (CDR3 | pMHC + V-region) show minimal residual effects. (C) Pairwise distance scale. Distributions of native whole-pose composite distances are shown for structure pairs grouped by receptor identity, peptide–MHC identity, and germline V-region sharing. The shaded region indicates the empirical reproducibility floor estimated from true recrystallization pairs (median ≈ 0.05). Structures sharing germline V-region framework cluster near this floor, whereas structures with different germline frameworks show substantially larger distances. (D) Localization of antigen-context effects. Adjusted values for MHC allele and peptide length contrasts are shown across selected FramePose blocks and axes. These effects localize primarily to orientation coordinates—particularly CDR3β and groove-normal twist axes—indicating that antigen context tunes docking geometry through local orientation adjustments rather than global receptor repositioning (* p < 0.10; I] FDR q < 0.05).

**Table 2.**
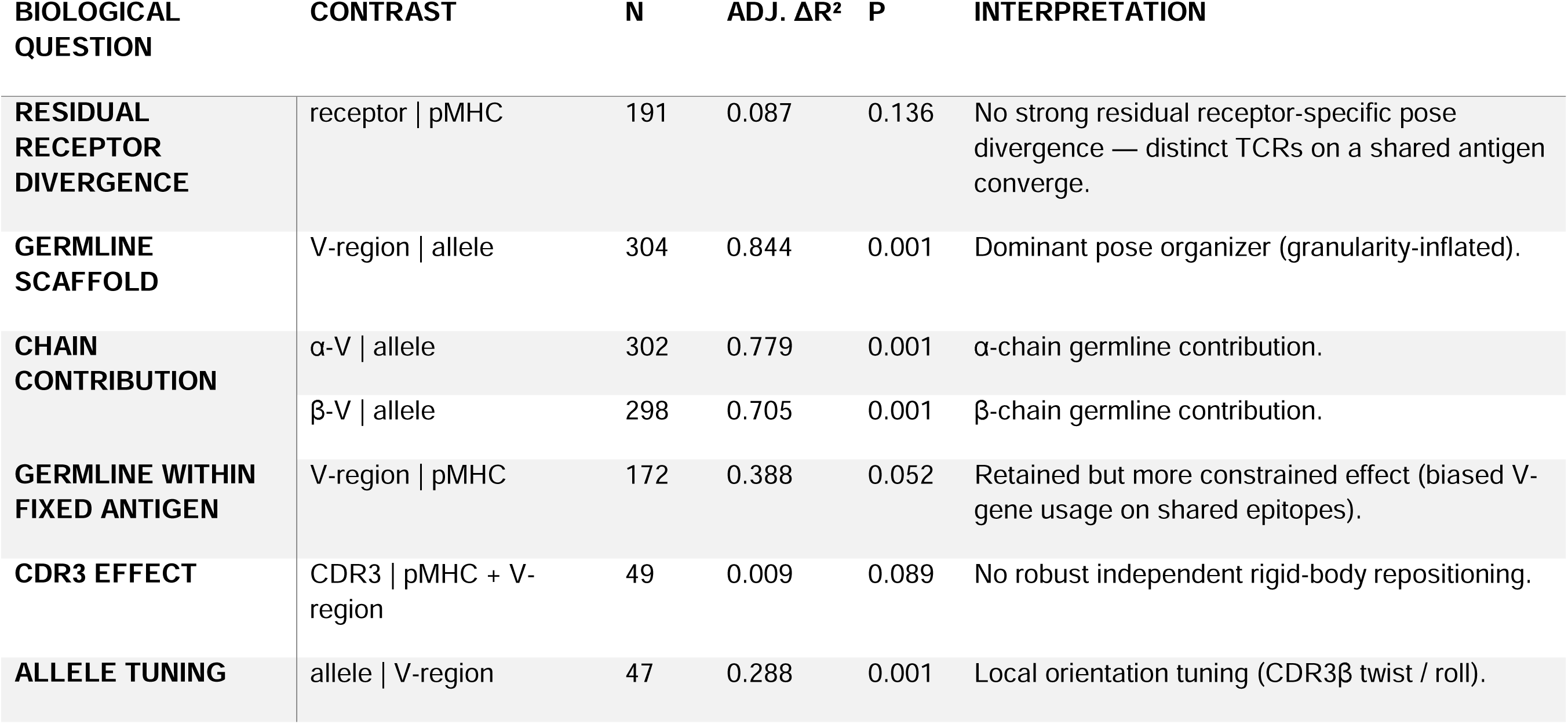
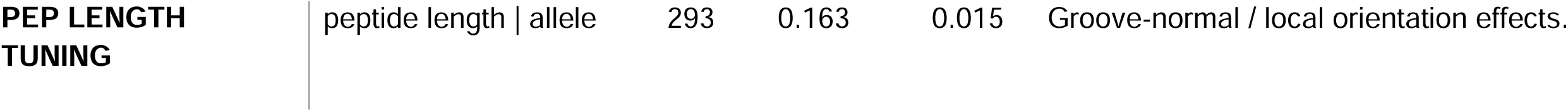
Conditioned partial PERMANOVA contrasts defining biological determinants of FramePose geometry. Each row represents a conditioned contrast of the form (*focal* | *stratum*), in which the contribution of a biological factor (focal) is evaluated after accounting for variation explained by one or more conditioning variables (stratum). This design isolates the independent association of sequence or antigen-context features with docking geometry while controlling for confounding structure in the dataset. For each contrast, the reported effect size is the adjusted Δ*R*^2^, defined as the additional fraction of distance-based variance explained by the focal factor beyond that explained by the conditioning variables. Higher Δ*R*^2^ values indicate stronger association of the focal factor with FramePose geometry within the specified context. All contrasts were evaluated using restricted permutation testing within conditioning strata (999 permutations), with Benjamini–Hochberg false-discovery-rate correction applied within contrast families. Only strata containing at least two levels of the focal factor were included to ensure identifiability of effects.

Consistent with this, pose-distance analyses showed that structures sharing germline V-region framework approach the reproducibility floor defined by recrystallized complexes, even when CDR3 sequences differ (Figure 5C; Supplementary Tables S4–S5). In contrast, structures with different germline frameworks are separated by substantially larger distances. These results indicate that TCRs recognizing the same pMHC tend to adopt shared docking geometries constrained by germline framework, rather than distinct receptor-specific poses.

We next identified the receptor features responsible for this shared geometry. Germline V-region identity was the dominant determinant of FramePose variation, showing that V-region identity explained the largest whole-pose effect when conditioned on MHC allele (V-region | allele, adjusted Δ*R*^2^ = 0.844, p = 0.001; Figure 5B; Table 2). Both α- and β-chain germline components contributed (Δ*R*^2^ = 0.779 and 0.705 for α-V | allele and β-V | allele, respectively; both p = 0.001), indicating that the shared receptor scaffold is not imposed by one chain alone but reflects paired germline architecture across the TCR variable domains.

These effect sizes should be interpreted with regards to the grouping granularity. Fine-grained labels with many small groups, such as V-region, receptor identity, α-V, and β-V, can produce large adjusted Δ*R*^2^ values partly because the data are partitioned into many sequence-specific groups. Therefore, their Δ*R*^2^ values should not be interpreted as calibrated fractions of variance directly comparable to coarser labels such as MHC allele or peptide length. Instead, the most informative comparison is between determinants with similar granularity and conditioning. In this like-for-like comparison, germline V-region retained a large whole-pose effect, whereas CDR3 sequence retained essentially none (Figure 5B; Table 2; Supplementary Figure S7A). This supports the germline V-region framework, rather than independent CDR3-sequence variation, as the main receptor-sequence feature associated with the FramePose rigid-body scaffold.

The germline framework remained the leading receptor-associated determinant even when the entire pMHC context was held fixed, although the effect was attenuated and borderline (V-region | pMHC, adjusted Δ*R*^2^ = 0.388, p = 0.052; Figure 5B; Table 2). This attenuation is expected if fixed antigens already constrain the V-regions represented in the cohort: a shared peptide–MHC target is recognized not by arbitrary receptor frameworks, but by a restricted set of germline solutions that impose similar receptor geometries.

Consistently, CDR3 sequence did not detectably reposition rigid-body pose after controlling for germline framework and pMHC context (CDR3 | pMHC + V-region, Δ*R*^2^ = 0.009, p = 0.089; Figure 5B; Table 2) and did not produce robust block-level effects across whole-TCR, CDR3α, or CDR3β reach, offset, or orientation (Supplementary Figure S7B–C). This result should not be read as evidence that CDR3 sequence is unimportant for recognition. This result indicates that CDR3 sequence does not independently drive rigid-body repositioning of the receptor, although it may influence recognition through mechanisms not captured by rigid-body frame representation. Because FramePose treats each CDR3 loop as a local rigid body, this analysis does not test internal CDR3 backbone conformation, side-chain geometry, or residue-level contact chemistry.

### 3.2 Antigen context tunes the germline scaffold through local orientation adjustments

We next examined how antigen context perturbs the germline-anchored docking scaffold. Compared with germline effects, MHC allele and peptide features showed smaller but structured contribution.

Fine allele variation within a fixed class and species was not significant (allele | class x species, adjusted Δ*R*^2^ = 0.03, p = 0.31; Supplementary Figure S7A), indicating limited pose variation at this level. However, within a fixed germline scaffold, allele identity produced a reproducible effect (allele | V-region, adjusted Δ*R*^2^ = 0.288, p = 0.001; Figure 5B; Table 2). Block- and axis-level decompositions localized this effect primarily to orientation coordinates, especially CDR3β and groove-normal axes (Figure 5D; Supplementary Figure S7B–C). These results indicate that allele variation tunes docking geometry within a shared germline scaffold rather than altering the scaffold itself.

Peptide length similarly contributed a smaller but significant effect (peptide length | allele, adjusted Δ*R*^2^ = 0.163, p = 0.015; Figure 5B; Table 2). This effect localized most clearly to local orientation coordinates rather than to a global receptor repositioning (Figure 5D; Supplementary Figure S7B). Coordinate-level decomposition further showed that this signal is concentrated in CDR3 coordinates, particularly along groove-normal twist axes (plen | allele, Δ*R*^2^ = 0.31 and 0.21 for CDR3α and CDR3β , respectively; Supplementary Figure S7C).

Peptide identity showed a detectable association in some conditioned analyses, but this association should be interpreted cautiously because peptide identity is not fully separable from the CDR3 sequence composition of the available structural cohort. We therefore treat peptide-identity effects as peptide-context associations rather than isolated peptide-sequence effects. Together, the allele and peptide-length results indicate that antigen context modulates the germline-defined receptor scaffold through localized FramePose adjustments, especially in CDR3β and groove-normal orientation axes.

Overall, we established a hierarchical model of TCR–pMHC docking organization. Germline V-region framework defines the primary docking scaffold, constraining receptor placement and orientation across structures. TCRs recognizing the same antigen therefore occupy similar docking geometries rather than exhibiting strong receptor-specific divergence. Antigen context, including MHC allele and peptide length, introduces smaller adjustments to this scaffold, primarily through local orientation changes, especially in CDR3β. In contrast, CDR3 sequence does not independently reposition rigid body pose once germline framework and antigen context are controlled. These results place the CDR3-local orientation association signals to BSA and binding affinity in a biological context: rather than reflecting arbitrary geometric variation, they likely correspond to antigen-dependent tuning of a germline-defined docking scaffold, particularly in CDR3-local orientation degrees of freedom.

### 4. CDR3β reach links FramePose geometry to interface burial and binding-associated structure

We previously showed that FramePose geometry contains nonredundant information associated with BSA and binding-affinity labels, but also that rigid-body pose alone provides only modest affinity association. Previous structural studies have established that TCR–pMHC recognition depends on multiple factors coupled, including docking geometry, interface burial, shape complementarity, and residue-level contacts, with CDR3 loops often providing local structural adjustments within a constrained receptor–MHC docking topology^17,33,35–37^. We therefore asked whether the pose-associated signals identified in the BSA and affinity association studies could be traced to a physically interpretable properties, specifically through CDR3-local relationship with interface burial, packing, or contact remodeling.

#### 4.1 Interface burial is the dominant cross-sectional correlate of affinity

Across the affinity-annotated cohort (n = 244), the strongest structural correlate of affinity was interface burial and packing, with buried surface area (BSA) providing the most consistent structural correlate. In mixed models with MHC allele as a random intercept, greater buried surface area (BSA) was associated with stronger binding (β = −0.414, p < 0.001; Figure 6A, Supplementary Table S6). Shape complementarity was weaker and only borderline (β = −0.193, p = 0.026; Supplementary Table S6).

**Figure 6.**
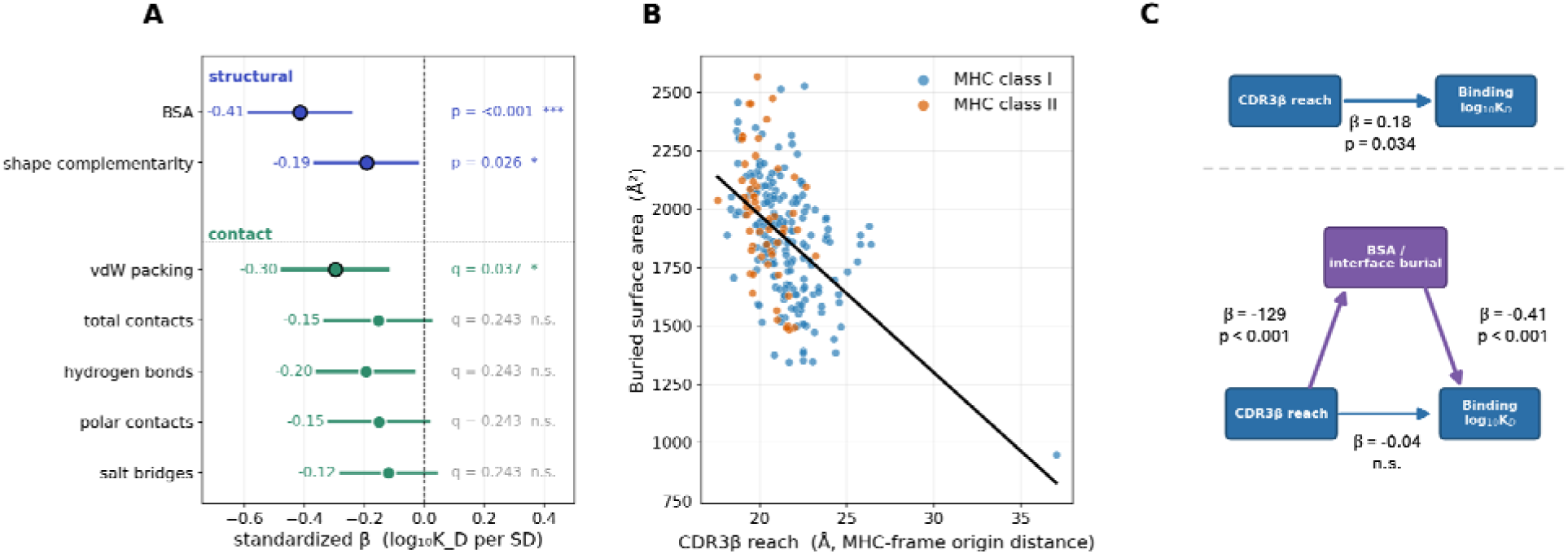
Interface burial mediates the relationship between CDR3β reach and binding affinity. Cross-sectional analyses were performed on n = 244 complexes with curated dissociation constants ( , μM). Associations were evaluated using linear mixed-effects models with MHC allele as a random intercept. (A) Interface burial and affinity. Cross-sectional association between buried surface area (BSA) and binding affinity ( ). Greater interface burial is associated with stronger binding (β < 0), indicating that interface size is the dominant structural correlate of affinity. Shape complementarity shows a weaker association (Supplementary Table S6). (B) CDR3β reach and interface burial. Association between CDR3β reach and BSA. Greater reach, corresponding to the CDR3β loop positioned farther from the pMHC groove, is associated with reduced burial. This relationship remains robust after allele-referencing and adjustment for germline V-region (Supplementary Figure S10), indicating that CDR3β reach captures within-allele geometric variation in interface size. (C) Mediation of reach–affinity association by BSA. Association between CDR3β reach and affinity before and after adjustment for BSA and shape complementarity. The reach–affinity association is modest and becomes non-significant after covariate adjustment, indicating that the effect of reach on affinity is mediated by interface burial rather than representing an independent pose-based contribution.

**Figure 7.**
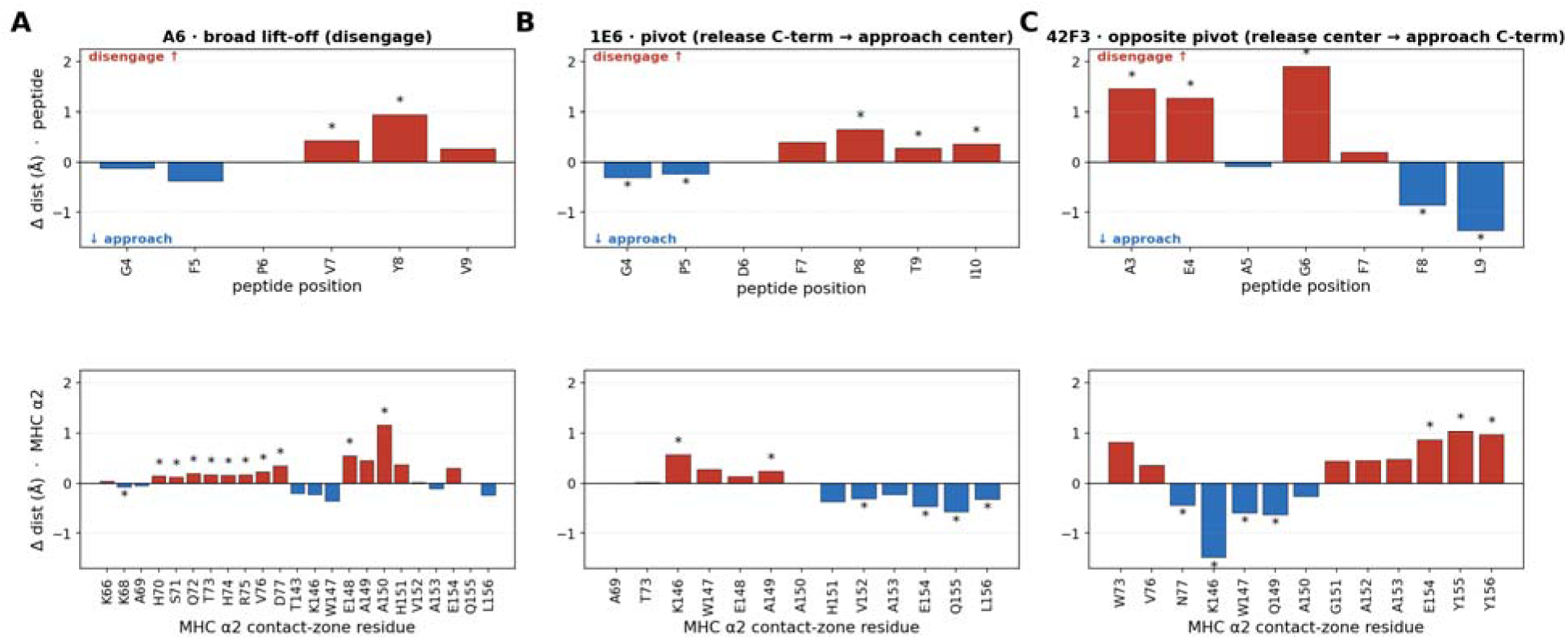
Residue-level CDR3β footprint remodeling in engineered peptide panels. Residue-level contact-zone analyses for selected engineered panels with sufficient size (n ≥ 5). Structural variation is quantified as differences in minimum Cα–Cα distances between CDR3β residues and peptide (top) or MHC contact-zone residues (bottom), relative to the strongest-binding structure within each panel. Each panel shows spatial maps of CDR3β–interface distances across peptide positions and adjacent MHC residues, highlighting regions of structural change associated with affinity variation. Positive values indicate increased distance (disengagement), and negative values indicate decreased distance (closer contact) (* : nominal Wilcoxon p < 0.05). (A) A6 panel. Affinity-weakening variants show broad CDR3β disengagement from both peptide residues (particularly C-terminal positions) and the MHC α2 helix, consistent with reduced interface engagement. (B) 1E6 panel. Variants show a pivot-like remodeling pattern, with increased distances at C-terminal peptide/MHC regions and decreased distances at central interface residues, indicating a redistribution of CDR3β contacts. (C) 42F3 panel. Remodeling occurs in the opposite direction relative to the 1E6 panel, with decreased distances at C-terminal peptide regions and increased distances at central positions, indicating a distinct receptor-specific shift in interface engagement. Across panels, structural changes localize to the CDR3β interface region, but the direction of remodeling varies between receptors. These patterns illustrate receptor-specific, context-dependent footprint adjustments rather than a single universal rule relating structural change to affinity perturbation.

Contact-based analyses supported the same interpretation. Among contact features, only van der Waals packing survived FDR correction (CDR3–peptide packing β = −0.297, BH-q = 0.023; CDR3–pMHC packing β = −0.296, BH-q = 0.023; Supplementary Figure S8A), whereas directional polar contacts such as hydrogen bonds and salt bridges were weaker and did not survive correction. However, when BSA and shape complementarity were included as covariates, this packing signal was no longer significant (CDR3–peptide packing β = −0.19, p = 0.04, BH-q = 0.24; CDR3–pMHC packing β = −0.01, p = 0.88; Supplementary Figure S8B), indicating that the packing–affinity association is carried by interface burial rather than acting as an independent mechanism.

#### 4.2 CDR3β reach provides a geometric readout of interface burial

We next asked which FramePose coordinate is most directly linked to interface burial. The clearest relationship is CDR3β reach, defined as the distance between the CDR3β local-frame origin from the pMHC groove frame. Greater CDR3β reach, corresponding to the CDR3β loop frame sitting farther from the groove, was strongly associated with reduced BSA (β = −129.2 Å² per SD of reach, p < 0.001; Figure 6B; Supplementary Figure S10A).

Because this association could reflect baseline differences among MHC alleles or TCR germline families rather than a direct geometric relationship, we tested two controls. First, we allele-referenced CDR3β reach by subtracting each allele’s mean reach, thereby removing between-allele differences in groove geometry and baseline burial. The reach–BSA relationship remained robust within alleles (β = −109 Å² per SD, p < 0.001; Supplementary Figure S10B,C). Second, because we have shown that the V gene identity can set the default CDR3β framework position over the groove, we included TRBV germline as a categorical covariate. The association again remained strong with only modest attenuation (β = −110.2 Å² per SD, p < 0.001; Supplementary Figure S10C,E). Thus, the reach–burial relationship is not explained simply by comparing different MHC alleles or different TRBV-defined docking baselines. These results indicate that CDR3β reach captures within-allele, germline-controlled variation in interface burial, rather than reflecting only differences across receptors or alleles.

CDR3β reach also showed a modest association with affinity (β =0.180, p=0.034; Supplementary Figure S9B). This effect was modest relative to the reach–BSA relationship. However, when BSA and shape complementarity were included as covariates, the direct reach coefficient was attenuated and no longer significant (β =-0.04, p=0.677; Figure 6C; Supplementary Figure S9C). This pattern indicates that the reach–affinity relationship is mediated by interface burial, rather than as an independent pose mechanism. Because reach and BSA are partly geometrically coupled — a loop frame sitting farther from the groove is expected to bury less surface — this should be interpreted as structural linkage, not as evidence for a causal pathway.

Not all pose coordinates reflect stable structural relationships. For example, the CDR3β cross-groove directional coordinate showed affinity association in the engineered-peptide subset and remained significant after adjustment for BSA and shape complementarity (Supplementary Figure S10F). However, this effect did not replicate in the full-cohort and weakened after allele referencing (Supplementary Figure S10F). Thus, directional pose–affinity associations appear to be cohort- or context-dependent, in contrast to the more robust reach–burial relationship.

#### 4.3 Within-panel analyses reveal system-specific remodeling

To test whether the reach–burial relationship predicts mutation-level effects, we analyzed engineered peptide panels in which receptor and MHC are fixed and peptide sequence varies. These panels consisted of sets of TCR–pMHC structures in which the receptor clonotype and MHC allele were held constant while the peptide sequence was experimentally varied, producing closely related complexes with different measured affinities. This design provides a stricter test than the cross-sectional cohort, as it removes most between-receptor, between-MHC, and between-germline associated variations that can drive population-level associations. Within each panel, we therefore asked whether contact remodeling or FramePose pose changes tracked affinity differences relative to the best-binding panel member.

Within panels, contact remodeling showed nominal trends but did not support a general pooled rule. Several CDR3β–peptide or CDR3β–pMHC hydrogen bond and polar contact features were nominally associated with affinity change in the expected direction, with contact loss corresponding to a larger affinity penalty (Supplementary Figure S11A). However, no contact-remodeling feature met both FDR and within-panel permutation criteria across the tested contact partitions (Supplementary Figure S11C). Thus, contact remodeling within engineered panels was suggestive in some systems but not stable enough to define a general mutation-level contact rule.

FramePose coordinates showed a similar trend, where panel-referenced pose changes were system-specific across mutant panels. CDR3β reach or cross-groove tilt showed nominal associations in some analyses (Supplementary Figure S11B), consistent with the cross-sectional reach–burial relationship, but these effects were not stable under the stricter panel-sensitive analyses (Supplementary Figure S11C). Thus, the absolute cross-sectional relationship between CDR3β reach and burial should not be read as a mutation-level rule that any increase or decrease in reach will predict Δ*K_D_* within a fixed receptor. Instead, it identifies a physically interpretable geometric coordinate whose population-level relationship to interface burial is clear, while within-panel affinity changes remain system-specific.

Residue-level footprint analysis illustrates why a universal within-panel model may be difficult. We selected the three best-powered engineered panels, A6, 1E6, and 42F3 (n≥ 5) to map mutation-associated CDR3β distance changes across peptide positions and nearby MHC contact-zone residues. CDR3β remodeling occurred in a similar broad structural region — the central-to-C-terminal peptide and adjacent MHC α2 helix — but the direction of remodeling differed by receptor. In A6, affinity-associated changes were accompanied by broader CDR3β lift-off from peptide and MHC residues. In 1E6, CDR3β followed a pivot-like pattern, releasing from C-terminal peptide/MHC region while approaching more central interface residues. In 42F3, the direction was reversed, the release from central peptide/MHC residues and approach toward the C-terminal peptide region. These results provide an illustrative structural vignette showing that CDR3β defines a recurrent local geometric arena for interface organization, but residue-level affinity responses are TCR-specific.

FramePose geometry to physical interface properties and clarifies the scope of pose–affinity relationships. Across complexes, affinity is primarily associated with interface burial, and CDR3β reach provides a geometric coordinate that tracks this relationship. However, the reach–affinity association is mediated by BSA and does not represent an independent effect of pose. More generally, rigid-body pose does not provide a universal affinity model. While cross-sectional geometric relationships are identifiable, mutation-level structural responses remain context-dependent. FramePose therefore contributes by identifying interpretable geometric coordinates—particularly CDR3β reach and orientation—that relate local pose to interface organization, rather than by replacing contact- or energy-based models of binding.

## Discussion

TCR–pMHC recognition is shaped by sequence, chemistry, and three dimensional geometry, yet commonly used geometric descriptors summarize docking predominantly at the whole-receptor level. In this study, we introduce TCR-FramePose as a local-frame representation that decomposes the pose of the whole TCR, CDR3α, and CDR3β relative to a common pMHC groove frame. By separating reach, offset direction, and orientation for each body, FramePose provides a structured framework for distinguishing global receptor placement from local CDR3-loop position and orientation. The central contribution of this work is to establish a coordinate system that makes these layers of TCR–pMHC geometry separable, interpretable, and quantitatively comparable across structures.

This decomposition is useful in two complementary ways. First, FramePose recovers established large-scale features of TCR–pMHC docking, including class-associated receptor placement and reverse-polarity outliers. Second, it exposes local orientation variation that is largely hidden from conventional global descriptors. This distinction is important because existing descriptors already capture much of the global placement signal, including receptor distance and centroid positioning over the pMHC platform. The added value of FramePose arises from CDR3-local geometry, particularly CDR3 orientation, which reveals an orientation layer of variation that conventional descriptors only weakly capture.

An apparent tension in our results is that the FramePose blocks contributing the most nonredundant information in cross-validated BSA and affinity analyses are CDR3-local orientation features, whereas the clearest allele-controlled interface interpretation arises from CDR3β reach. We interpret this difference as reflecting the distinct questions addressed by the two analyses rather than a contradiction. The predictive analyses assess which FramePose components provide information beyond conventional docking descriptors. In this setting, CDR3β orientation was the most consistent FramePose-specific contributor, with additional signal from CDR3α orientation and CDR3β offset. This pattern is expected if conventional descriptors already encode much of the global distance and positional information but do not capture how CDR3 loops are locally rotated over the peptide–MHC surface. We therefore interpret the CDR3β orientation signal as reflecting local footprint organization - how the β loop is oriented over peptide and MHC contact zones and therefore which interface regions are approached or released - rather than simply how close the loop lies to the groove.

The mixed-effects interface analyses address a different question by explicitly conditioning on MHC allele and testing which geometric features retain stable relationships to interface burial. This relationship can be viewed as a physical calibration: a loop frame positioned farther from the groove necessarily buries less surface. The attenuation of the reach–affinity association after adjustment for BSA and shape complementarity supports the interpretation that reach is primarily reflects burial-linked geometry rather than an independent determinant of binding affinity. Thus, CDR3β orientation and CDR3β reach describe complementary aspects of loop engagement. Orientation captures a context-sensitive, FramePose-specific footprint organization, whereas reach provides an allele-controlled measure of burial depth.

These observations clarify the biological interpretation of the two signals. CDR3β orientation represents the primary source of nonredundant geometric information captured by FramePose, reflecting local rotational positioning that is not encoded in conventional features. In contrast, CDR3β reach provides a direct link to interface size and burial under allele-controlled models. These coordinates therefore distinguish between local footprint orientation and overall burial depth, offering a more nuanced description of CDR3β engagement. Importantly, this distinction reinforces a central conclusion of the study: FramePose captures a meaningful structural layer of TCR–pMHC recognition, but rigid-body pose alone does not constitute a complete affinity mode.

The biological grouping analyses place this local geometric layer within a broader hierarchy of TCR–pMHC organization. Across the structural cohort, germline V-region framework is the dominant determinant of rigid-body docking geometry, whereas CDR3 sequence did not detectably reposition this geometry once antigen context and germline framework are controlled. This observation does not imply that the CDR3 sequence is unimportant for recognition. Instead, it indicates that the rigid-body pose captured by FramePose is largely scaffolded by the variable-domain framework, while CDR3 sequence likely contributes through mechanisms not fully represented in a rigid-loop frame, including internal backbone flexibility, side-chain interactions, peptide contacts, and energetic optimization.

MHC allele and peptide length contributed smaller but informative perturbations to this scaffold. Their effects localize primarily to CDR3β and groove-normal orientation axes, consistent with antigen-dependent tuning of local loop geometry rather than wholesale rearrangement of the receptor pose. This interpretation provides context for the CDR3β orientation signals observed in predictive analyses; they may reflect allele- or antigen-dependent adjustments to loop positioning. When allele baselines are explicitly modeled in the interface analyses, part of this structured variation is absorbed by the allele term, whereas the more general reach–burial relationship remains detectable. Thus, conditioning on allele shifts the question from identifying nonredundant geometric features to identifying stable geometric correlates of interface structure.

The engineered panel analyses further delineate the limits of this generalization. Within fixed TCR–MHC panels, peptide mutations do not produce a universal rule for contact remodeling or pose difference associated with affinity changes. Instead, residue-level footprint analyses reveal a recurrent structural region centered on the CDR3β interface, but with receptor-specific directions of remodeling. This finding argues against a simple model in which a single geometric coordinate predicts affinity changes across systems. Rather, FramePose identifies where structural variability tends to occur—often within the CDR3β footprint—while the functional consequences of that variability depend on receptor-specific and context-dependent factors.

Several limitations follow this interpretation. The structural cohort is not a random sample of TCR–pMHC recognition, and It is enriched for crystallized complexes, engineered variants, and well-studied antidens. Receptors, peptides, and MHC alleles are also unevenly represented and partially confounded, and although partial PERMANOVA and mixed-effects modeling mitigate these effects, they do not fully eliminate them. Similarly, the BSA and binding affinity association models rely on repeated random cross-validation, which is optimistic because related complexes can appear across training and test folds. Accordingly, these models should be interpreted as cross-validated structural association analyses rather than as deployment-ready predictors of affinity. In addition, affinity measurements are heterogeneous and unevenly distributed, and binarization of KD values simplifies evaluation at the cost of resolution.

FramePose also has methodological limitations. It represents the TCR and CDR3 loops as rigid local frames and therefore does not capture side-chain chemistry, loop flexibility, peptide conformational changes, solvent effects, or energetic contributions. The framework also depends on deterministic frame definitions; although these were validated with geometric diagnostics, alternative frame constructions—such as contact-weighted or peptide-centered frames—may capture additional aspects of recognition. These limitations do not undermine the utility of the framework but define its scope as a geometric abstraction rather than a full physical model.

Practically, FramePose provides a standardized coordinate framework for structure-guided TCR analysis and downstream design workflows. In the context of in silico TCR–pMHC modeling, including structure prediction and docking pipelines, FramePose enables systematic screening of predicted complexes by quantifying whether candidate poses fall within physically and biologically plausible regions of receptor placement and CDR3-local geometry. This allows large sets of predicted TCR–pMHC models to be filtered or ranked based not only on global docking metrics but also on CDR3-local orientation and positioning, which are poorly captured by conventional descriptors. In TCR engineering applications, FramePose coordinates can be used to monitor how sequence perturbations — particularly in CDR3 regions — propagate to changes in loop placement, orientation, and interface engagement, providing a geometric readout complementary to energetic or contact-based scoring. More broadly, the decomposition into reach, offset, and orientation offers a compact set of interpretable features that can be integrated into machine-learning or design pipelines to guide model selection, constrain docking solutions, or prioritize variants with favorable geometric properties. These uses do not make FramePose a standalone predictor of affinity or specificity, but position it as a useful geometric layer for organizing structural models and linking sequence perturbations to interpretable changes in receptor pose.

In summary, TCR-FramePose extends conventional docking analysis by decomposing pose into reach, offset direction, and orientation across both whole-TCR and CDR3-local frames. This representation supports a model in which germline V-region framework defines a shared docking scaffold, antigen context tunes that scaffold through localized orientation adjustments, and functional variation often manifests in CDR3-local geometry. FramePose does not replace sequence-, contact-, or energy-based analyses but complements them by providing a standardized geometric layer for interpreting TCR–pMHC recognition. More broadly, these results support the view that TCR–pMHC docking is best understood not as a single global pose, but as a hierarchy of coupled global and local geometric degrees of freedom.

## Data and Code Availability

All structural data are publicly available through TCR3d^22^ (https://tcr3d.ibbr.umd.edu) and the Protein Data Bank^38^ (https://www.rcsb.org). The TCR_CoM^14^ tool is available at https://github.com/EsamTolba/TCR-CoM. The TCRdock^16^ software implementing the SE(3) parameterization of Bradley et al. is available at https://github.com/phbradley/TCRdock. Euler angle extraction code and processed data are deposited at https://github.com/KChen-lab/TCR-FramePose.

## Author Contributions

K.H.K. performed the majority of data analysis and interpretation and drafted the manuscript. X.J., Q.Y., V.M., M.D., and K.C. revised the manuscript and contributed to data interpretation. K.H.K., A.R., and K.C. conceived and supervised the project, guided the analytical framework, and finalized the manuscript. All authors reviewed and approved the final version of the manuscript.

## Acknowledgments

The authors thank Yejin Kim from UTHealth Houston School of Biomedical Informatics, Khaled Sanber and Maura L. Gillison from MD Anderson Cancer Center for many helpful discussions.

