## Supplementary Figure for "TCR-FramePose: a local-frame representation for decomposing global docking and CDR3 loop geometry in TCR-pMHC recognition"

**Supplementary Figure S1. Frame-construction diagnostics for pMHC, TCR, and CDR3 frames.** Per-body frame stability diagnostics are shown for whole-TCR, CDR3α, and CDR3β frames. Singular-value decomposition (SVD) summaries quantify axis stability, with x- and y-axes representing principal-axis variance ratios. Additional diagnostics include CDR3 loop-shape descriptors and apex geometry metrics, including apex position and apex dominance. These diagnostics confirm orthogonality, right-handedness, and numerical stability of all constructed frames, supporting the robustness of the FramePose representation across the structural cohort.

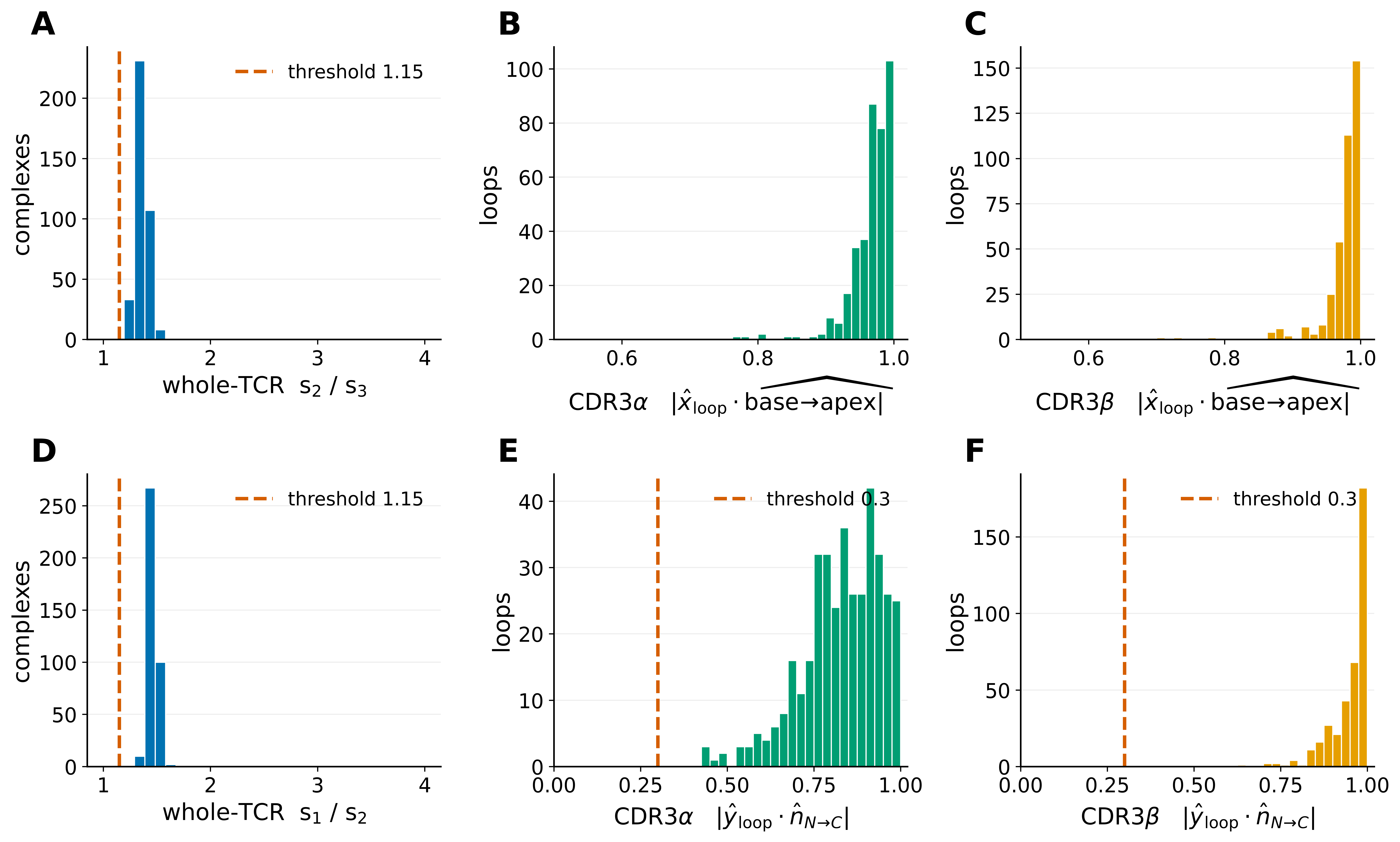

**Supplementary Figure S2. Distance-correlation structure of FramePose blocks.**

Pairwise distance correlation between the nine FramePose blocks (three bodies × three manifolds) computed across n = 378 TCR–pMHC complexes. For each block, pairwise distances were defined in native geometry: absolute Euclidean differences for reach, geodesic distances on $S^{2}$ for offset direction, and quaternion geodesic distances on $SO(3)$ for orientation.

Distance matrices were double-centered before computing distance correlation. Warmer colors indicate stronger dependence. Strong coupling is observed among corresponding offset and orientation blocks across bodies, reflecting coordinated global and CDR3-local docking geometry, whereas reach shows weaker coupling with directional and rotational components.

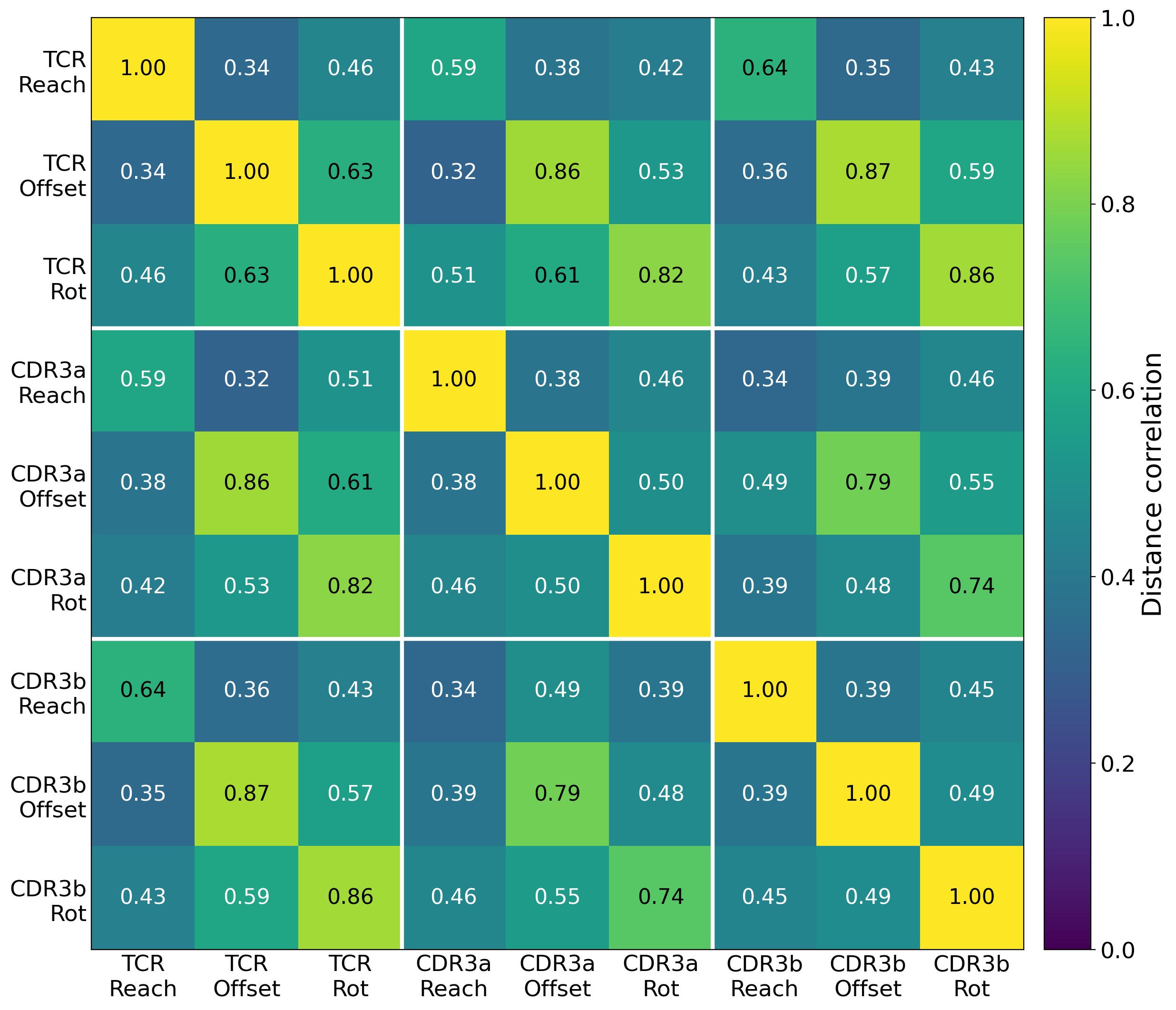

**Supplementary Figure S3. Principal component structure of FramePose tangent coordinates.**

Principal component analysis of the 18-dimensional FramePose tangent representation across n = 378 TCR–pMHC complexes (282 class I, 96 class II). Prior to analysis, coordinates were centered and scaled within each block to ensure balanced contributions across reach, offset, and orientation components.

(A) Explained variance (per-PC and cumulative) and class separation measured by Cliff’s $\delta$.

(B) Class separation in the PC1–PC4 plane (top class-discriminating PCs). Ellipses indicate $2\sigma$ covariance.

(C) Block-level contributions computed as the sum of squared coordinate loadings per block.

(D) Top contributing tangent coordinates to class-discriminating PCs. This analysis identifies dominant modes of variation and highlights the contribution of specific FramePose components to class separation.

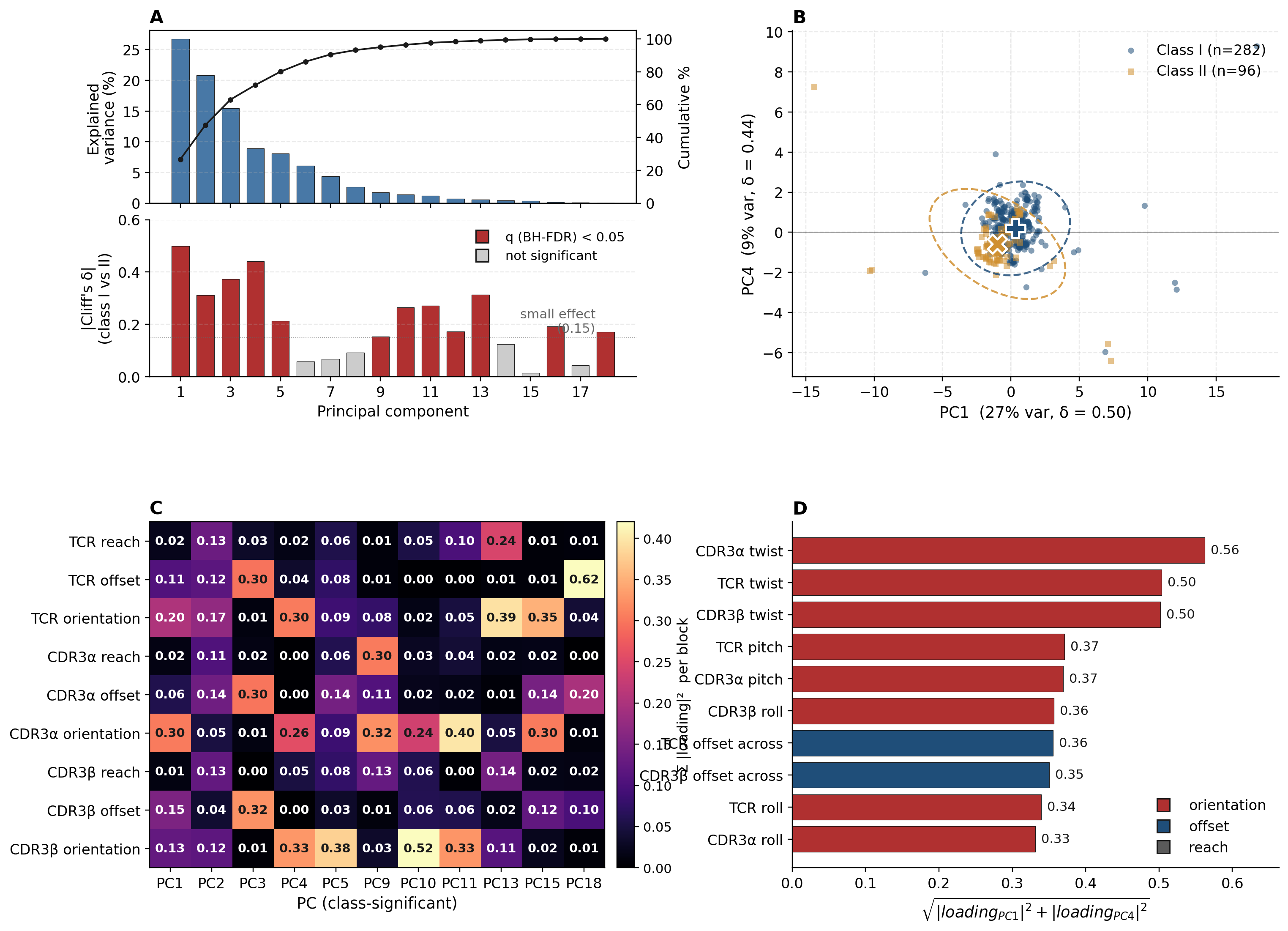

**Supplementary Figure S4. Conventional descriptors detect polarity outliers but do not resolve orientation sub-modes.**

Canonical reverse-polarity and related outliers are visualized in conventional descriptor space using crossing angle and TCR-CoM $\phi$.

(A) Distribution of crossing angle across the cohort. Many polarity outliers show elevated crossing angles; however, some outliers (e.g., 9RUP) fall within the main distribution, indicating incomplete detection.

(B) Crossing angle versus TCR-CoM $\phi$. Canonical reverse-polarity structures split into class-dependent clusters due to centroid-position differences rather than true orientation differences. FramePose resolves these structures as a unified orientation mode by representing rotations directly on $SO(3)$, separating rotational effects from translational placement.

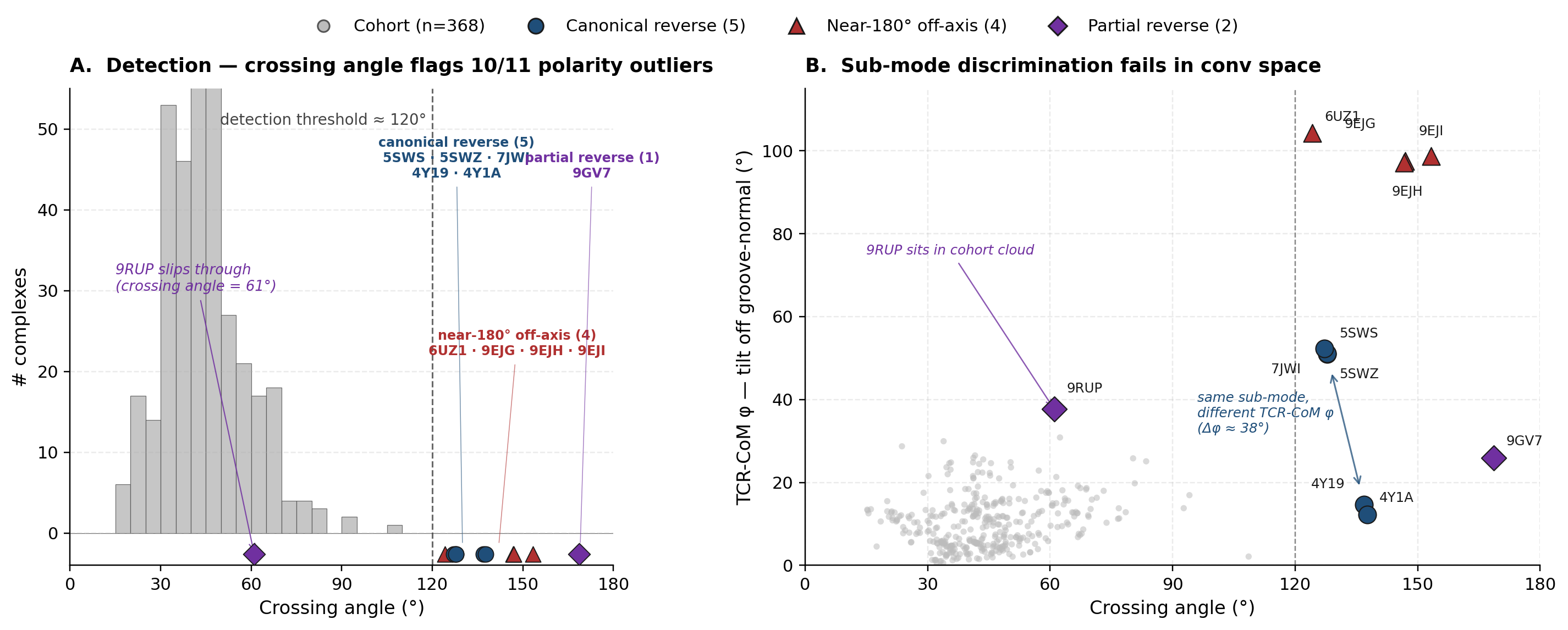

**Supplementary Figure S5. Reverse augmentation analysis of conventional descriptors added to saturated FramePose models.**

Each conventional docking descriptor set was added individually to the saturated FramePose model for the corresponding task to evaluate whether conventional descriptors provide additional association signal beyond FramePose. Bars show the change in association strength relative to the saturated FramePose baseline (Δmetric), where positive values indicate improvement after inclusion of the conventional descriptor. Error bars represent empirical 2.5–97.5 percentile intervals across 20 repeated cross-validation splits.

(A–B) BSA regression. Conventional descriptors were added to the saturated all-bodies FramePose model. Changes in association strength are shown for coefficient of determination (ΔR²; A) and Spearman rank correlation (Δρ; B).

(C–D) Affinity classification. Conventional descriptors were added to the saturated CDR3α+CDR3β FramePose model. Changes in association strength are shown for AUROC (ΔAUROC; C) and AUPRC (ΔAUPRC; D).

Minimal or non-significant changes indicate that conventional descriptors provide little additional association signal beyond FramePose, supporting the conclusion that FramePose captures the majority of geometry-associated information represented in conventional docking descriptors.

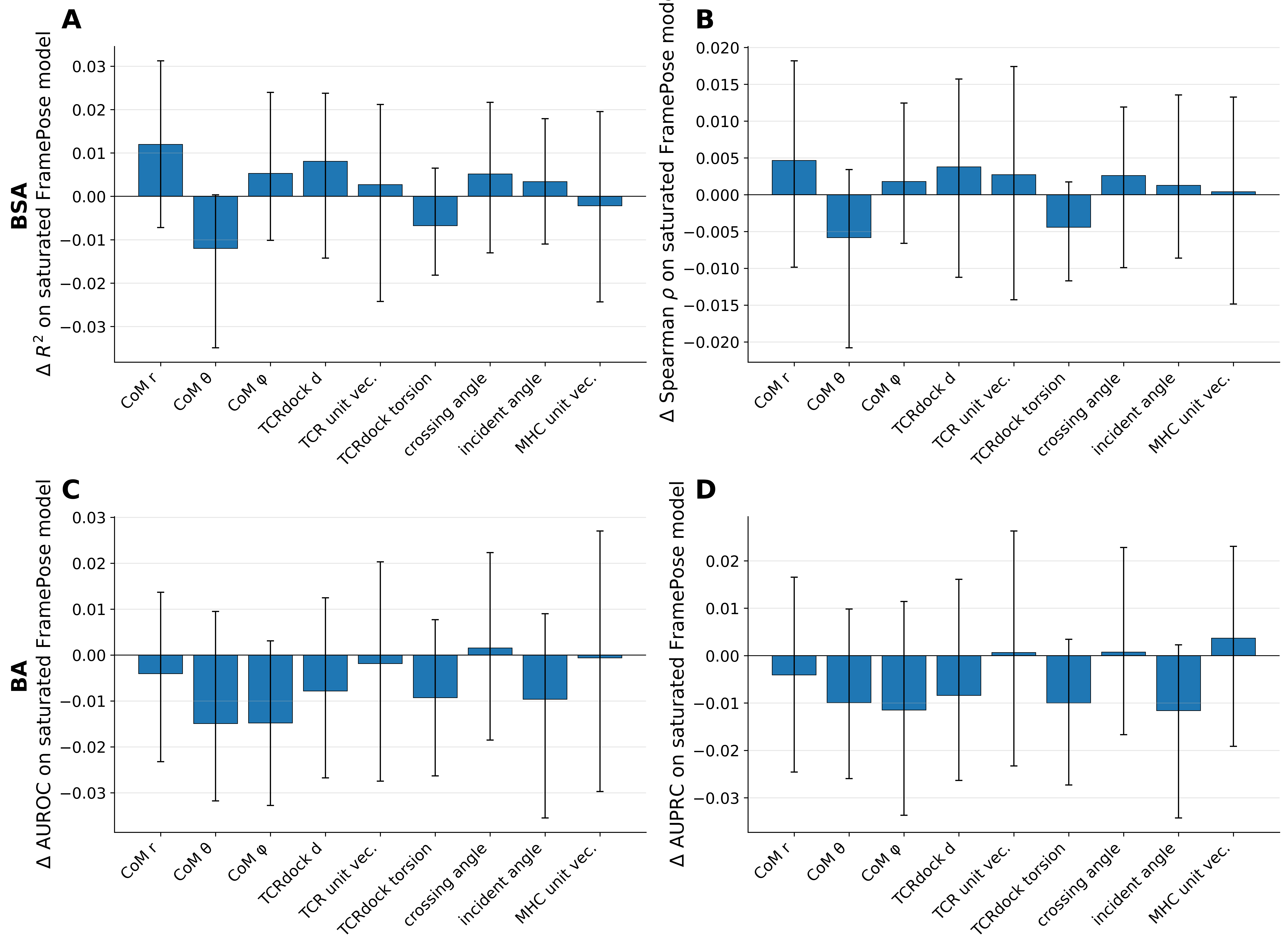

**Supplementary Figure S6. Recoverability of FramePose features from conventional docking descriptors.**

Recoverability quantifies how well FramePose geometric features can be reconstructed from conventional descriptor sets. FramePose variables were predicted from the 16-dimensional conventional descriptor set using cross-validated regression models. Bars show mean out-of-fold $R^{2}$, and error bars represent empirical 2.5–97.5 percentile intervals across repeated cross-validation splits. The dashed line indicates $R^{2}$ = 0.5 for reference.

(A) Block-level recoverability. Each bar represents the joint recoverability of a FramePose block defined by body (whole TCR, CDR3α, or CDR3β) and feature type (reach, offset, or orientation). Scalar reach values were evaluated directly, while offset and orientation blocks were evaluated using multicomponent targets. Labels above bars indicate residual errors (Å for reach; degrees for offset and orientation). Whole-TCR reach shows the highest recoverability, whereas CDR3 orientation blocks show the lowest recoverability.

(B) Per-coordinate recoverability. Each bar represents the $R^{2}$ for one of the 18 FramePose tangent coordinates. Conventional descriptors more effectively reconstruct global placement features (reach and offset) than local orientation coordinates, particularly for CDR3 loops.

Lower recoverability indicates geometric variation not captured by conventional descriptor families. These poorly recoverable features correspond to those contributing most strongly to association models (Figure 3-4), highlighting CDR3-local orientation as the primary source of nonredundant geometric information in FramePose.

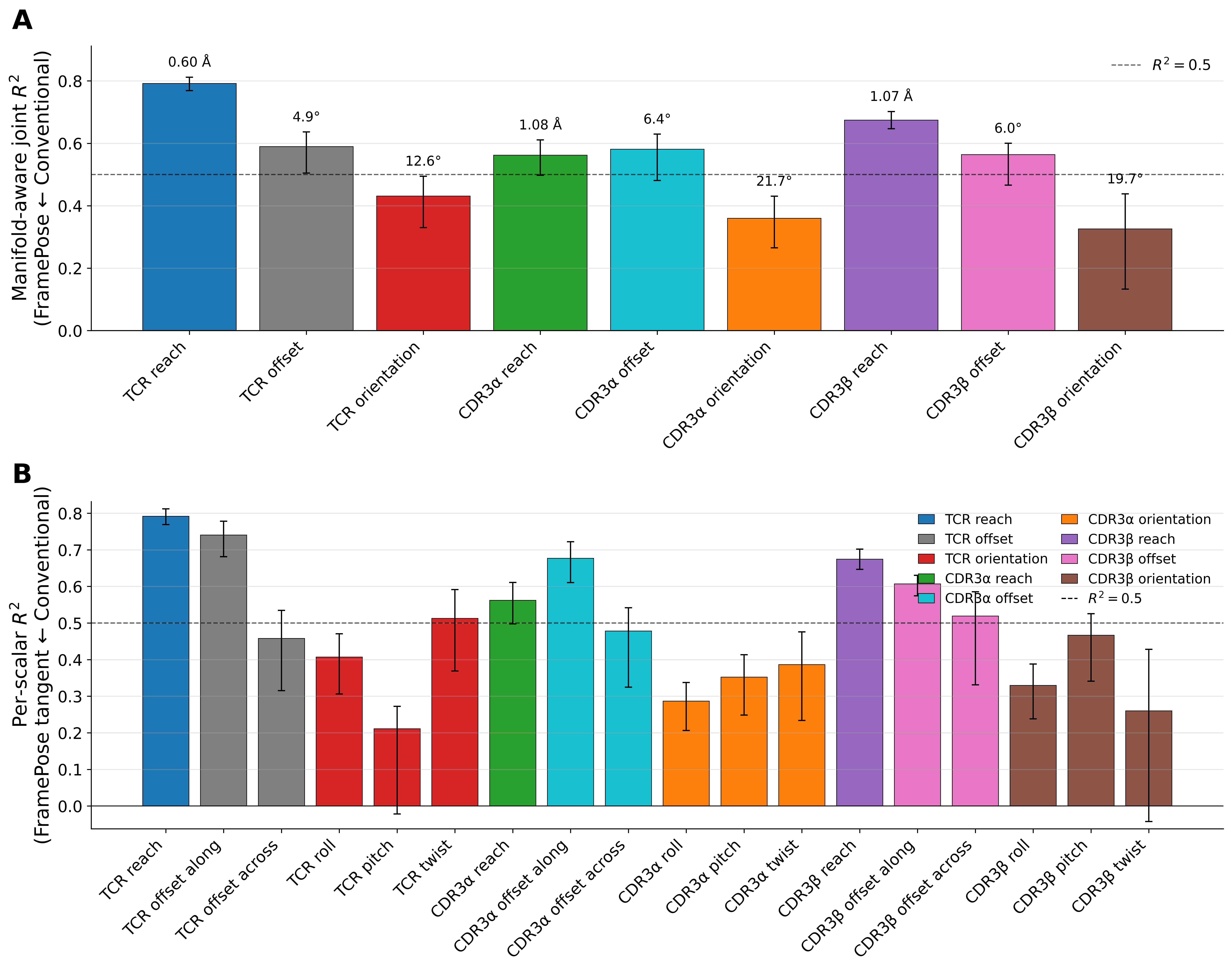

**Supplementary Figure S7. Full decomposition of conditioned partial PERMANOVA analyses for biological determinants of FramePose geometry.**

All analyses were performed on n = 378 αβTCR–pMHC complexes unless otherwise noted. Biological determinants were evaluated using Type-II partial PERMANOVA under conditioned contrast designs of the form (focal factor | conditioning variables), with restricted permutations performed within conditioning strata (999 permutations). Effect sizes are reported as adjusted $\Delta R^{2}$.

(A) Whole-pose determinant effects. Adjusted $\Delta R^{2}$ values are shown for all tested contrasts at the level of the whole-pose composite distance. Points represent effect sizes, and annotations indicate the number of eligible structures, permutation p-values, and FDR-corrected significance. Germline V-region identity shows the strongest association with docking geometry, whereas contrasts involving CDR3 sequence show minimal residual effects after conditioning. Antigen-context determinants (MHC allele and peptide length) show smaller but significant contributions.

(B) Block-level localization. Block-level decomposition of each determinant across the nine FramePose blocks (whole TCR, CDR3α, CDR3β × reach, offset, orientation). Adjusted $\Delta R^{2}$ values indicate which geometric components are associated with each determinant. Germline V-region effects are distributed across global and local pose components, whereas antigen-context effects localize predominantly to orientation blocks, particularly within the CDR3β frame. CDR3 sequence shows minimal contribution across all blocks when conditioned on germline and antigen context.

(C) Axis-level (coordinate) localization. Axis-level decomposition across the 18 tangent coordinates. Each panel reports adjusted $\Delta R^{2}$ values for individual axes, allowing localization to specific geometric directions. Antigen-context effects (allele and peptide length) are concentrated in CDR3-local orientation coordinates, particularly along groove-normal (twist) axes, whereas reach and offset coordinates show minimal contributions. Germline V-region effects extend across multiple axes, reflecting its role in defining the overall docking scaffold.

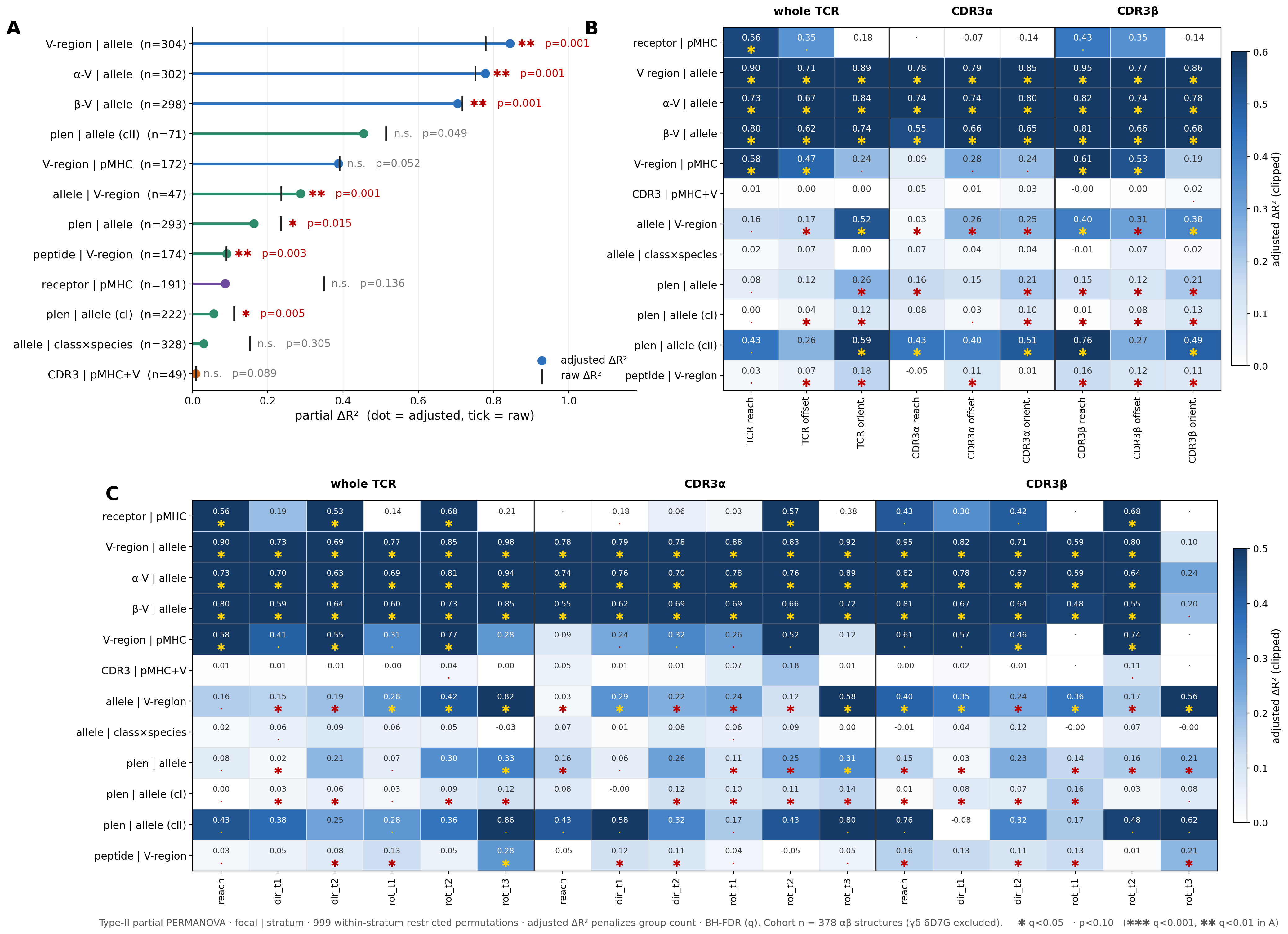
Across levels of analysis, these results support a hierarchical organization of docking geometry: germline V-region framework defines the primary docking scaffold, while antigen-context variables (MHC allele and peptide length) introduce smaller, localized adjustments in CDR3 orientation, particularly along groove-normal axes. CDR3 sequence does not independently reposition rigid-body pose after conditioning on germline framework and antigen context.

**Supplementary Figure S8. Contact-feature associations with binding affinity are largely explained by interface burial.**

Analyses were performed on n = 244 affinity-annotated TCR–pMHC complexes using linear mixed-effects models with MHC allele as a random intercept. Contact features were evaluated across loop sets, interface targets (peptide or pMHC), and interaction types, with predictors $z$-standardized prior to model fitting. Points show fixed-effect coefficients and horizontal bars show 95% confidence intervals. Printed values indicate nominal p-values ( * : p < 0.05, ★: BH-q < 0.05).

(A) Unadjusted contact–affinity associations. Cross-sectional associations between contact features and binding affinity ($\log_{10} K_{D}$). Several features, particularly van der Waals packing interactions involving CDR3 loops, show significant associations after FDR correction, with increased packing associated with stronger binding.

(B) Adjusted models with interface covariates. The same analyses after including buried surface area (BSA) and shape complementarity (SC) as covariates. Contact-feature associations are substantially attenuated and no longer significant after multiple-testing correction, indicating that contact-level signals largely reflect interface burial rather than independent contributions.

These results support the interpretation that affinity-associated contact features are mediated by interface size and packing.

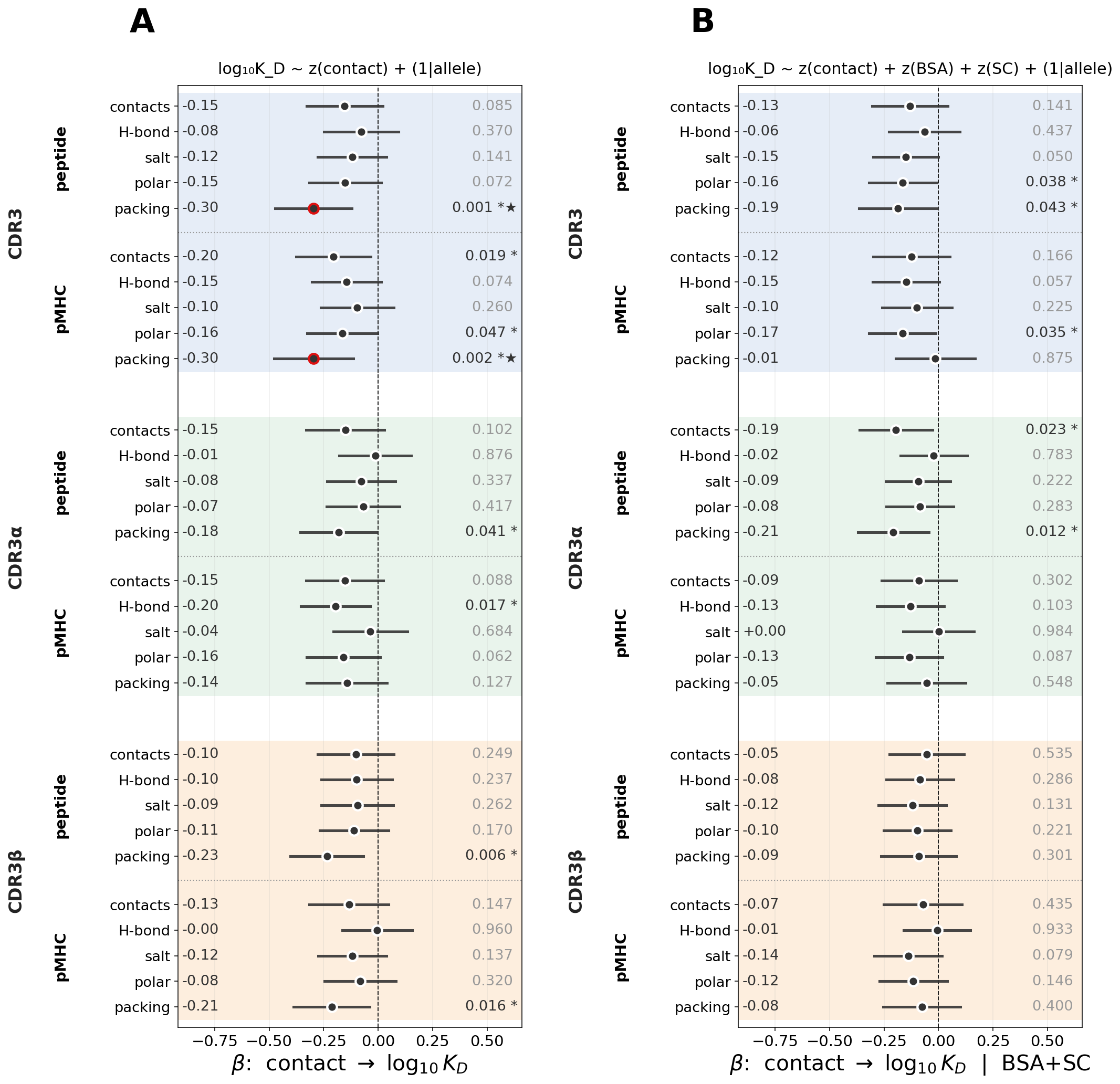

**Supplementary Figure S9. Cross-sectional associations between FramePose coordinates and binding affinity.**

Analyses were performed on n = 244 complexes using linear mixed-effects models with MHC allele as a random intercept. Predictors were z-standardized, so $\beta$ represents the response change per one standard deviation of the coordinate. Points show fixed-effect coefficients and horizontal bars show 95% confidence intervals. Printed values indicate nominal p-values (* : p < 0.05, ★: BH-q < 0.05).

(A) Coordinate-level associations. Estimated effects (β) for individual FramePose tangent coordinates on $\log_{10} K_{D}$. Most coordinates show weak or non-significant associations, consistent with the modest overall predictive signal observed in classification analyses (Figure 4).

(B) CDR3β reach–affinity association. Association between CDR3β reach and affinity, showing a modest effect in which larger reach corresponds to weaker binding.

(C) Adjusted reach–affinity model. The reach–affinity association after including BSA and shape complementarity as covariates. The reach effect is attenuated and becomes non-significant, indicating mediation by interface burial.

Together, these results show that most pose–affinity associations are weak and that the CDR3β reach signal is largely explained by its relationship with interface burial.

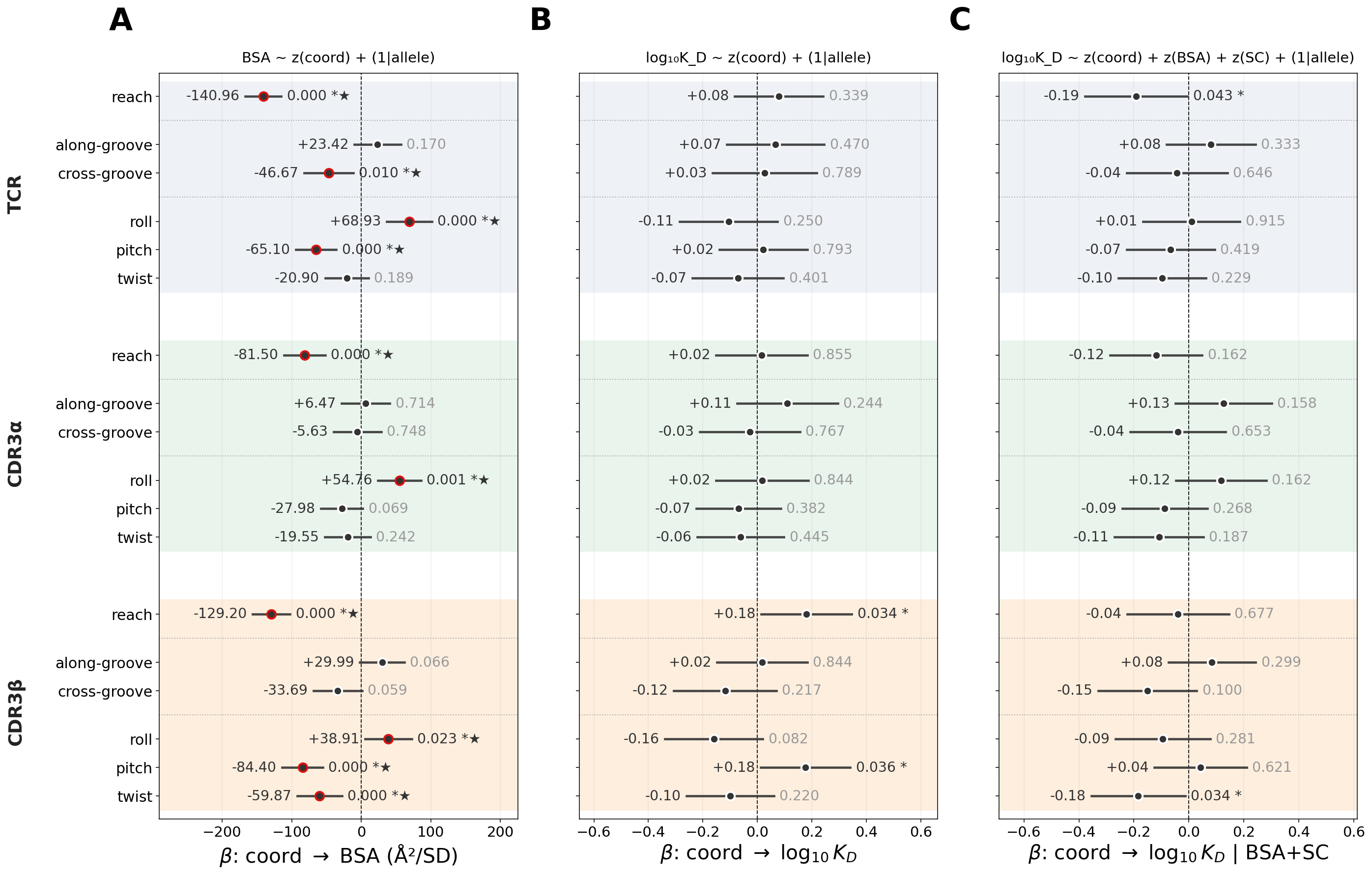

**Supplementary Figure S10. Robustness of the CDR3β reach–burial relationship and sensitivity of CDR3β cross-groove.**

Analyses were performed using linear mixed-effects models with MHC allele as a random intercept unless otherwise noted.

(A) Baseline reach–BSA association. Cross-sectional relationship between CDR3β reach and buried surface area (BSA).

(B) Allele-referenced CDR3β reach versus BSA. CDR3β reach was centered within each MHC allele by subtracting the allele-specific mean prior to standardization, thereby isolating within-allele variation. The reach–burial association remained strong, indicating that the relationship is not explained by between-allele differences in groove geometry or baseline docking depth.

(C) Stability of reach–BSA coefficients across robustness models. Estimated coefficients for the reach–BSA relationship across multiple model specifications, including the raw model, a TRBV-adjusted model, and the allele-referenced model. The negative association remained consistent across all models, demonstrating that the reach–burial relationship is robust to both germline and allele-level adjustments.

(D) Class-stratified reach–BSA relationship. The negative association between CDR3β reach and BSA is observed independently within both MHC class I and class II subsets, supporting a shared geometric interpretation across MHC classes.

(E) Added-variable (residual) analysis controlling for allele and TRBV. BSA residuals are plotted against reach residuals after regressing out MHC allele and TRBV germline effects. The relationship persists under this stricter control, although attenuated, indicating that CDR3β reach retains within-context information about interface burial beyond these covariates.

(F) Sensitivity analysis for a CDR3β directional coordinate.

Association between a representative CDR3β cross-groove orientation coordinate and binding affinity ($\log_{10} K_{D}$). This coordinate shows an association in the engineered-peptide subset and remains significant after adjustment for BSA and shape complementarity, but the effect weakens after allele-referencing and does not replicate in the full cohort. This contrast highlights that directional pose–affinity associations are cohort- or context-dependent, whereas the CDR3β reach–burial relationship (panels B–E) is robust.

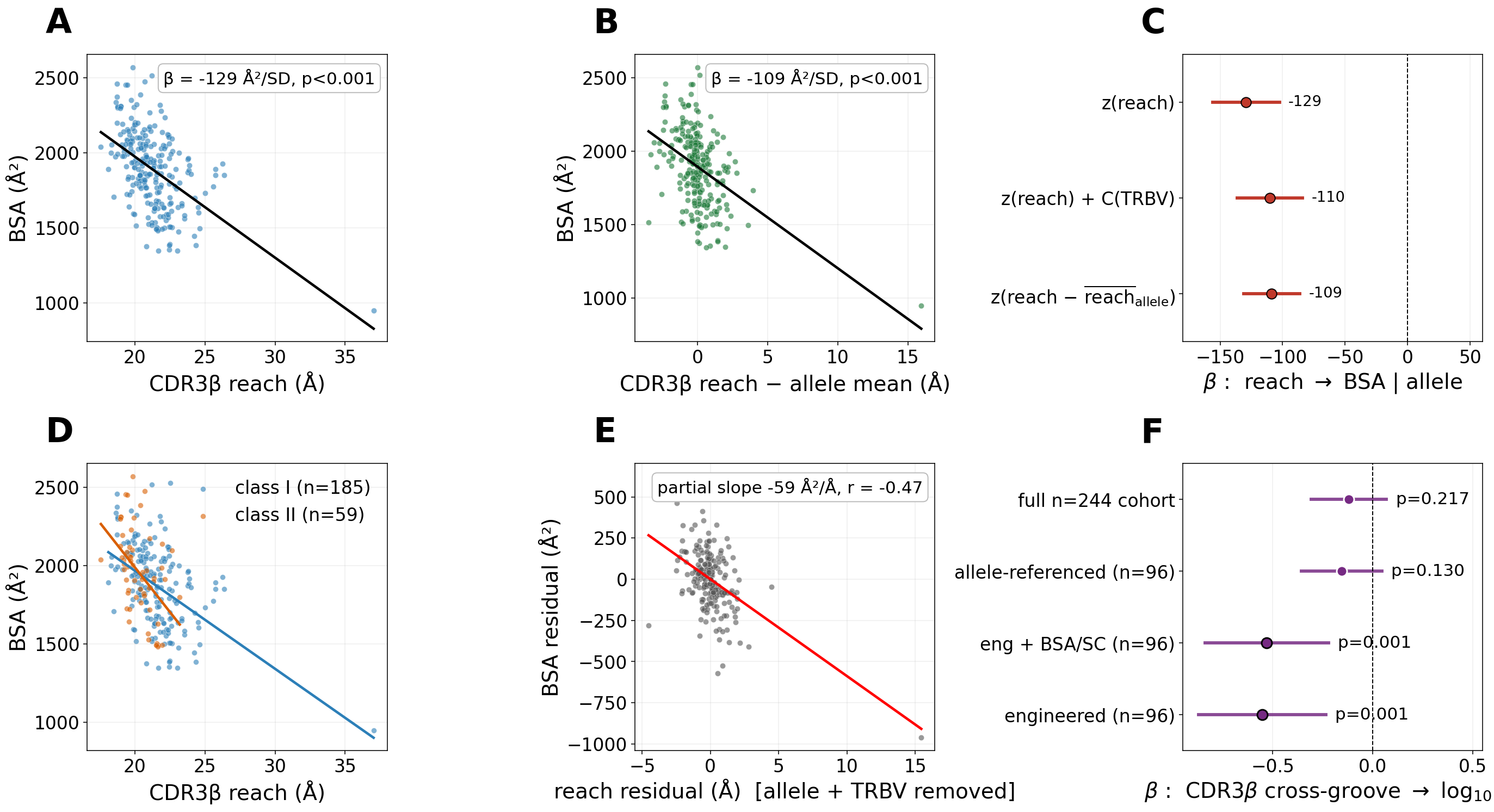

**Supplementary Figure S11. Within-panel evaluation of contact remodeling and FramePose drift in engineered peptide panels.**

Analyses were performed on engineered peptide panels (n = 96 complexes across 37 panels), in which each complex was compared to the strongest-binding reference structure from the same panel. Binding changes were quantified as ΔΔG, and structural remodeling was represented as panel-referenced changes in contact features or FramePose coordinates.

(A) Contact-remodeling associations. Estimated coefficients for panel-referenced changes in contact features across CDR3 loop sets and peptide or pMHC targets. Negative coefficients indicate that loss of the corresponding contact feature is associated with a larger affinity penalty.

(B) FramePose drift associations. Estimated coefficients for panel-referenced changes in FramePose coordinates. Nominal associations are observed for CDR3β reach and selected directional coordinates, including CDR3β cross-groove offset and whole-TCR cross-groove offset.

(C) Significance summary across feature families. Bars show the number of features reaching nominal significance (p < 0.05), FDR significance (BH q < 0.05), and within-panel permutation significance (p < 0.05). Permutation testing was applied to the top-ranked features within each family. No contact-remodeling or FramePose-drift feature satisfies both FDR and permutation criteria, indicating that mutation-level affinity variation does not collapse into a single pooled structural rule.

(D) Panel-size distribution. Distribution of structures per panel in the engineered-peptide cohort. Most panels contain only two structures, with relatively few larger panels. This limited and unbalanced sampling motivates the use of panel-random-intercept models and within-panel permutation testing.

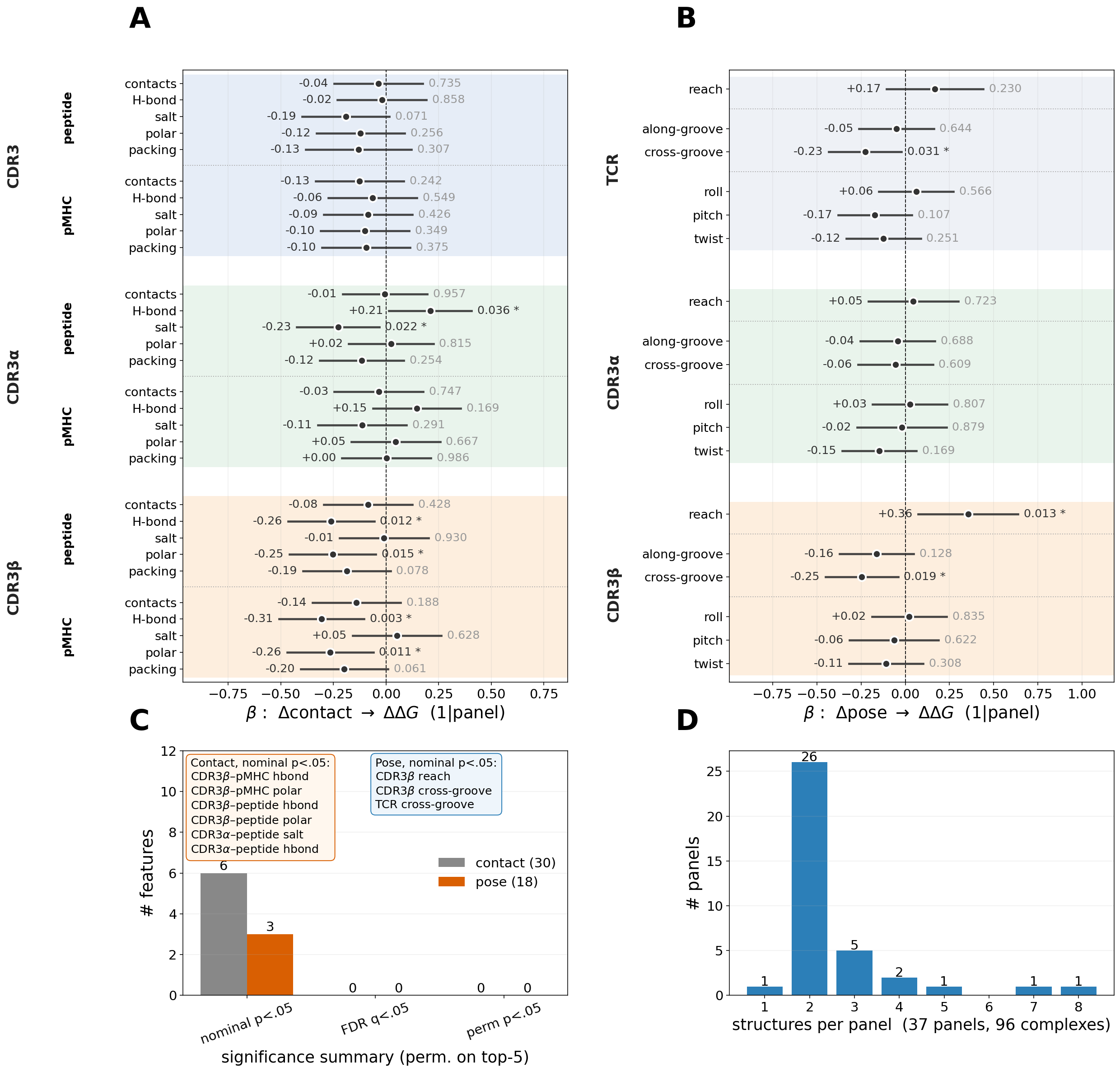

**Supplementary Table S1. Comparison of TCR–pMHC docking descriptor families.**

Conventional and FramePose descriptor families are compared in terms of geometric representation, dimensionality, and interpretability. Columns indicate the native parameters, the level of representation (whole receptor versus local loops), whether translation and orientation are explicitly separated, the underlying manifold structure, and interpretability with respect to pMHC-aligned axes.

TCR-CoM descriptors represent receptor placement using spherical coordinates of the TCR centroid relative to the pMHC frame. Crossing and incident angles describe projected aspects of docking but do not span full rigid-body degrees of freedom. TCRdock provides a compact six-parameter representation of whole-receptor transformations. FramePose extends these approaches by explicitly decomposing translation (reach and offset) and orientation for both the whole receptor and CDR3-local frames within a unified coordinate system.

| Descriptor | Native parameters | Whole vs local loop | Translation / orientation separation | Native manifold | Interpretable axes | Intended use |
| --- | --- | --- | --- | --- | --- | --- |
| TCR-CoM^1^ | r, θ, φ (3 DOF) | Whole receptor (centroid) | Translation only; no orientation | r ∈ ℝ⁺,  (θ, φ) ∈ $S^{2}$ | Centering & direction over groove; no orientation | Where the receptor sits over pMHC |
| Crossing^2^ / incident angle^3^ | crossing, incident (2 DOF) | Whole receptor | Partial; projected S^1^-like angular observables | No (projected scalars) | One docking angle each; not pMHC-fixed | Gross docking diagonal / tilt |
| TCRdock^4^ | d,  pMHC→TCR unit dir,  TCR→pMHC unit dir,  torsion $\tau$ (6 DOF) | Whole receptor | Yes, but orientation split across reciprocal direction + torsion | $d\in$ ℝ⁺,  $\hat{u}_{pMHC\to TCR}\in S^{2}$,  $\hat{u}_{TCR\to pMHC}\in S^{2}$,  $\tau\in S^{1}$ | Limited; not fixed roll/pitch/twist | Whole-TCR docking comparison; template-guided modeling |
| FramePose | 3 components per body (6 DOF per body) | Whole TCR + CDR3α + CDR3β | Explicit per body: reach + offset vs orientation | $reach\in\mathbb{R}^{+}$,  $offset\in S^{2}$, $orient\in SO(3)$  with tangent map | Roll / pitch / twist + reach + offset, pMHC-aligned | Decomposed global & local pose; outlier & polarity detection; localizing interface-associated variation |

**Supplementary Table S2.** **Noncanonical TCR–pMHC docking outliers identified by whole-TCR** $\boldsymbol{SO(3)}$ **orientation.**

Complexes are ranked by whole-TCR orientation deviation, defined as the $SO(3)$ geodesic distance (degrees) between each complex’s orientation and the cohort Fréchet-mean orientation. Axis rot denotes the groove-normal (twist) component of the tangent-space orientation vector, and axis rot fraction denotes the fraction of total rotation aligned with this axis. Off-axis rotation is defined as the combined contribution of groove-axis (roll) and cross-groove (pitch) components. CDR3α and CDR3β orientation deviations are shown for comparison. Strong correlation with whole-TCR orientation indicates that outlier signal reflects global receptor reorientation rather than isolated loop distortion. Outliers are categorized as canonical reverse polarity, near-180° off-axis flip, partial-reverse geometry, or substantial tilt based on rotation magnitude and axis alignment.

| PDB | TCR rot (°) | Axis rot (°) | Axis rot fraction | Off-axis rot (°) | CDR3α rot (°) | CDR3β rot (°) | Category |
| --- | --- | --- | --- | --- | --- | --- | --- |
| 9EJI | 179.1 | -101.9 | 0.57 | 147.2 | 174.9 | 172.6 | Near-180° off-axis flip |
| 9EJG | 177.9 | 104.7 | 0.59 | 143.8 | 176.1 | 173.2 | Near-180° off-axis flip |
| 9EJH | 176.2 | 106.7 | 0.61 | 140.2 | 176.9 | 172.9 | Near-180° off-axis flip |
| 5SWS | 172.5 | 165.2 | 0.96 | 49.8 | 177.4 | 179.9 | Canonical reverse polarity |
| 7JWI | 172 | 162.4 | 0.94 | 56.5 | 171.7 | 178.2 | Canonical reverse polarity |
| 4Y19 | 171.9 | 164.6 | 0.96 | 49.4 | 177.1 | 175.5 | Canonical reverse polarity |
| 5SWZ | 171.5 | 163.1 | 0.95 | 53.2 | 173.7 | 178.8 | Canonical reverse polarity |
| 4Y1A | 170.2 | 162.3 | 0.95 | 51.3 | 175.7 | 177.2 | Canonical reverse polarity |
| 6UZ1 | 166.9 | 128.6 | 0.77 | 106.3 | 171.7 | 161.3 | Near-180° off-axis flip |
| 9GV7 | 134 | 132.2 | 0.99 | 21.7 | 144 | 134.2 | Partial-reverse |
| 9RUP | 110 | -90.2 | 0.82 | 62.8 | 124.4 | 105.2 | Substantial tilt |

**Supplementary Table S3. Definition of biological grouping variables and conditioned contrasts used in Section 3 partial PERMANOVA analyses.**

Grouping variables used in Section 3 were derived from sequence-resolved TCR and pMHC annotations and tested using conditioned partial PERMANOVA designs. This table defines the sequence-derived and antigen-context grouping variables used to construct conditioned contrasts of the form (focal factor | conditioning variables). Each grouping variable is derived from IMGT/ANARCI-annotated TCR regions or curated TCR3d metadata. These definitions establish the conceptual framework for interpreting adjusted $\Delta R^{2}$ values reported in Table 2 and Supplementary Figure S7.

| Grouping / contrast | Definition | Fields used | Conditioning | Interpretation |
| --- | --- | --- | --- | --- |
| V-region identity | Paired germline framework label | TCRα/TCRβ FR1–FR3 + CDR1/CDR2, excluding FR4 | allele or pMHC | tests germline scaffold |
| Receptor identity | Full clonotype label | V-region + CDR3α/CDR3β | pMHC | tests residual receptor-specific pose divergence |
| CDR3 sequence | Paired junctional loop sequence | CDR3α + CDR3β | pMHC + V-region | tests independent CDR3 rigid-body repositioning |
| MHC allele | Resolved allele | sequence-resolved MHC allele | V-region or class$\times$species | tests allelic tuning |
| Peptide length | peptide residue count | peptide sequence length | allele | tests length-dependent groove accommodation |

**Supplementary Table S4. Recrystallization groups used to estimate the FramePose reproducibility floor.**

This table lists curated groups of structures representing true recrystallization pairs, defined as complexes with identical receptor, peptide, and MHC sequences and no engineered mutations or ambiguous residues. Each group contains multiple structures corresponding to nominally identical TCR–pMHC complexes.

For each group, pairwise native whole-pose FramePose distances were computed, and summary statistics (median and maximum distances) are reported. The pooled median distance across all audited recrystallization pairs (~0.05) provides an empirical estimate of the reproducibility floor, representing the minimum structural variation expected from crystallographic and construct-level differences.

This reproducibility floor serves as a reference scale for interpreting pairwise distance distributions and PERMANOVA effect sizes in Section 3, particularly when assessing whether observed pose differences reflect biological determinants or are within expected structural variability.

| Group | PDB IDs | Receptor | Peptide | MHC | n | Median dist. | Max dist. |
| --- | --- | --- | --- | --- | --- | --- | --- |
| R1 | 1OGA, 2VLJ, 2VLK | JM22 | GILGFVFTL | HLA-A*02:01 | 3 | 0.143 | 0.195 |
| R2 | 3D39, 3D3V, 3QFJ | A6 | LLFGFPVYV | HLA-A*02:01 | 3 | 0.033 | 0.038 |
| R3 | 7NME, 7NMF | 4C6 | QLPRLFPLL | HLA-A*24 | 2 | 0.039 | 0.039 |
| R4 | 7PBC, 7PDW | c728; c796 | GLYDGMEHL | HLA-A*02:01 | 2 | 0.065 | 0.065 |
| R5 | 8RYM, 8RYO | S2; S2-198 | ELFSYLIEK | HLA-A*03 | 2 | 0.033 | 0.033 |
| R6 | 6MKR, 6MNO | 5287; 6235 | RVSYYGPKTSPVQ | I-Ab | 2 | 0.134 | 0.134 |

**Supplementary Table S5. Pairwise FramePose distance scale for receptor, germline, and antigen-context relationships.**

This table summarizes native whole-pose FramePose distances across all admissible pairs of structures (n = 378), stratified by shared or differing biological features, including receptor identity, peptide–MHC identity, germline V-region framework, and CDR3 sequence.

Pair categories are defined to reflect increasing biological divergence, ranging from nominally identical complexes (same receptor and peptide–MHC) to structurally unrelated pairs (different germline V-regions). For each category, summary statistics (median, interquartile range, mean, and standard deviation) are reported.

Distances are additionally scaled relative to the reproducibility floor estimated from recrystallization pairs (Supplementary Table S4), enabling comparison of biologically meaningful pose differences to baseline structural variability.

These distance summaries provide an empirical scale for interpreting Section 3 PERMANOVA results, showing that structures sharing germline V-region framework cluster near the reproducibility floor, CDR3 sequence variation produces relatively small pose differences within a shared framework, and major pose divergence occurs primarily between distinct germline frameworks.

| Pair category | n | Median | IQR | Mean | SD |  | Interpretation |
| --- | --- | --- | --- | --- | --- | --- | --- |
| recrystallization (reproducibility floor) | 10 | 0.052 | 0.0346–0.1196 | 0.078 | 0.056 | 1.0$\times$ | empirical lower bound; structural reproducibility |
| Same receptor & same pMHC (nominal same complex) | 38 | 0.064 | 0.0424–0.1116 | 0.164 | 0.536 | 1.2$\times$ | includes recrystallizations and engineered same-complex series |
| Same germline V-region, same pMHC, diff CDR3 | 40 | 0.132 | 0.0741–0.2669 | 0.176 | 0.119 | 2.5$\times$ | CDR3 alone shifts pose only slightly. |
| Same germline V-region, diff pMHC | 226 | 0.141 | 0.0781–0.3066 | 0.245 | 0.252 | 2.7$\times$ | antigen-context tuning on a shared scaffold |
| Diff germline V-region, same pMHC | 214 | 0.413 | 0.2411–0.7111 | 0.653 | 0.958 | 7.9$\times$ | distinct germline frameworks give distinct poses |
| Diff germline V-region | 70949 | 0.800 | 0.6386–1.0203 | 1.003 | 0.816 | 15.4$\times$ | germline framework is the dominant organizer |

**Supplementary Table S6. Mixed-effects models of binding affinity as a function of interface burial and shape complementarity.**

Buried surface area (BSA) and shape complementarity (SC) were each evaluated as predictors of binding affinity using separate linear mixed-effects model applied to the cross-sectional affinity cohort (n = 244 structures spanning 41 MHC alleles). Predictors were z-standardized prior to model fitting, and MHC allele was included as a random intercept to account for shared structural context. Fixed-effect coefficients *(*$\beta$) are reported per one standard deviation increase in the predictor, together with 95% confidence intervals, standard errors, Wald $z$-statistics, and p-values. Negative coefficients indicate association with lower $K_{D}$, corresponding to stronger binding.

BSA shows a strong and highly significant association with affinity, whereas SC shows a weaker, marginal association. These results support interface burial as the dominant structural correlate of binding affinity in the cross-sectional cohort.

| Predictor | $\boldsymbol{\beta}$ | 95% CI | SE | z | p |
| --- | --- | --- | --- | --- | --- |
| BSA | −0.414 | [−0.583, −0.246] | 0.086 | −4.82 | 1.4×10⁻⁶ |
