## Supplementary Note for "TCR-FramePose: a local-frame representation for decomposing global docking and CDR3 loop geometry in TCR-pMHC recognition"

### Supplementary Note 1. Dataset curation

TCR3d crystal structures of αβTCR–pMHC complexes were the source of all structural data. After removing one structure (6D7G), identified as a $\gamma\delta$TCR, the cohort comprised 378 complexes (282 class I, 96 class II). TCRα/TCRβ chain identities and CDR annotations were taken from the curated TCR3d metadata and verified against the corresponding PDB chains; the pMHC platform was defined by the class I heavy chain or by both class II α and β chains. Of the 378 complexes, 377 had all conventional descriptors present (7BYD failed TCRdock parameterization and was excluded from the conventional-comparison cohort). Only Cα atoms from standard amino-acid residues were used to provide a consistent, noise-robust representation of backbone geometry across structures while avoiding dependence on side-chain modeling artifacts. Non-standard residues, modified residues, and heteroatoms (ligands, ions, waters) were excluded.

### Supplementary Note 2. pMHC, TCR, and CDR3 frame construction

#### FramePose coordinates were constructed in two stages. First, native rigid-body frames were defined for the pMHC reference, the whole TCR, and the CDR3α and CDR3β loops. Second, relative pose descriptors were computed between each body frame and the pMHC reference frame and exported as translation vectors, orientation matrices, quaternions, and accompanying frame-quality diagnostics. The procedures below describe frame definitions, orientation conventions, and stability criteria.

#### 2.1 pMHC reference frame

The pMHC reference frame was constructed from the peptide-binding platform using C$\alpha$ coordinates. For class I, the platform corresponded to the MHC class I heavy-chain peptide-binding domain (residues 1–180). For class II complexes, the platform included the MHC class II α1 (residues 1–90) and β1 (residues 1–95) domains.

Principal axes of the platform were defined using singular-value decomposition (SVD) of the centered Cα coordinate cloud yielding three orthonormal axes and three ordered singular values, $s_{1}\geq s_{2}\geq s_{3}$. The dominant SVD axis ($s_{1}$) was taken as the groove long axis, and the smallest-variance axis ($s_{3}$) was taken as the groove normal. These axes were then oriented using biologically anchored sign conventions.

The groove long axis was oriented to align with the peptide N→C direction, such that the vector connecting the first and last peptide Cα atoms had positive projection onto the long axis. The groove normal was oriented to point toward the TCR-facing side, such that the vector from the platform centroid to the TCR Cα centroid had positive projection onto the normal. The third axis was defined by the right-hand rule, producing a right-handed orthonormal frame with axes:

- $x_{\mathrm{MHC}}$: along the groove
- $y_{\mathrm{MHC}}$: across the groove
- $z_{\mathrm{MHC}}$: normal to the groove

The origin of the pMHC frame was defined as the centroid of the peptide-binding platform.

Frame quality and stability were assessed by evaluating orthogonality (pairwise axis dot products), right-handedness (determinant of the frame matrix), and sign-anchor consistency. The ratio $s_{1}$/$s_{2}$ was recorded as a long-axis stability diagnostic. Structures with near-degenerate first two axes (i.e., $s_{1}$/$s_{2}$ ≤1.15) were excluded to ensure a well-defined groove orientation.

#### 2.2 Whole-TCR frame

The whole-TCR frame was constructed from the combined Cα coordinates of the TCRα and TCRβ variable domains. Coordinates were centered at the TCR centroid, and principal axes were determined using SVD. The dominant principal axis $s_{1}$ was used as the approximate TCR body long axis. This axis typically captures the major elongation of the paired TCR variable domains.

To define the $\alpha$–$\beta$ orientation, the centroid difference between the TCR$\alpha$ and TCR$\beta$ domains was projected onto the plane perpendicular to the long axis ($s_{1}$). This projected vector defined the $\alpha\to\beta$ axis. The third axis was obtained by the right-hand rule, yielding a complete orthonormal whole-TCR body frame.

Axis orientations were fixed using biologically meaningful directions. The long axis ($s_{1}$) was oriented from the TCR centroid toward the pMHC centroid, corresponding to the docking direction. The α→β axis was oriented from the TCRβ centroid toward the TCRα centroid, establishing a consistent convention across complexes. The third axis was determined by the right-hand rule.

Frame diagnostics included singular values and their ratios, orthogonality checks, determinant validation, and verification that sign-anchor projections were sufficiently large to ensure stable orientation.

#### 2.3 CDR3 loop frames

Frames for CDR3α and CDR3β loops were constructed independently based on loop geometry. CDR3 residue ranges were defined using IMGT/ANARCI numbering.

For each loop, the first and last Cα atoms defined the loop base chord. The midpoint of this chord was used as the frame origin. The loop apex was defined as the residue whose Cα atom had the greatest perpendicular distance from the base chord, representing the most protruding point of the loop.

The vector from the base-chord midpoint to the apex defined the loop protrusion direction, which typically points toward the interaction interface. SVD was applied to the centered loop coordinates, and the principal axis most aligned with the base$\to$apex vector was selected as the primary axis and oriented toward the apex.

To define the secondary axis, the loop N→C direction (base chord) was projected into the plane perpendicular to the protrusion axis and orthogonalized using Gram–Schmidt. The third axis was obtained via the right-hand rule. This construction ensures that each CDR3 frame is anchored at the loop base, oriented toward the loop apex, and aligned with the loop backbone direction.

#### Additional descriptors were recorded to quantify frame stability, including the apex residue identity, apex distance from the base chord, and the apex dominance ratio. The apex dominance ratio was defined as the ratio of the apex distance to the next-largest distance among CDR3 residues and provides a measure of how uniquely defined the apex is.

#### 2.4 Frame diagnostics and quality control

For all frames (pMHC, whole TCR, CDR3α, and CDR3β), a consistent set of diagnostics was recorded to assess geometric stability and reproducibility. These included:

- SVD singular values $s_{1}$, $s_{2}$, and $s_{3}$
- singular-value ratios $s_{1}$/$s_{2}$ and $s_{2}$/$s_{3}$
- pairwise axis dot products to assess orthogonality
- the determinant of the frame matrix to verify right-handedness
- the magnitude of the sign-anchor projections used to orient each axis.

For CDR3 frames, additional diagnostics included apex dominance and residual errors from Gram–Schmidt orthogonalization.

Quality thresholds were applied to ensure well-defined frames across the curated cohort:

- Singular-value ratios had to exceed the prespecified stability thresholds
- Pairwise axis dot products < ${10}^{-6}$
- Determinants were within ${10}^{-9}$ tolerance of +1
- CDR3 apex dominance > 1.1
- Sign-anchor projections > ${10}^{-4}$

These criteria ensured that all retained frames were geometrically stable and consistently oriented. Diagnostic summaries were recorded for all structures to verify reproducibility and to support downstream analyses.

### Supplementary Note 3. Tangent-space geometry

For each body - whole-TCR, CDR3$\alpha$, or CDR3β - the pose relative to the pMHC frame is represented as a rigid-body transformation consisting of a translational vector and a relative orientation matrix,

$$T_{MHC\leftarrow body}=\left[ \begin{matrix} R_{rel} & t_{rel} \\ 0 & 1 \end{matrix} \right].$$

The relative translation vector is defined as the body-frame origin expressed in pMHC coordinate system,

$$t_{rel}=R_{MHC}^{T}\left( o_{body}-o_{MHC} \right),$$

and the relative orientation matrix is

$$R_{\mathrm{rel}}=R_{\mathrm{MHC}}^{T}R_{body}.$$

Thus, $t_{rel}$ describes the position of the body relative to the pMHC groove axes, and $R_{\mathrm{rel}}$ describes its orientation within the same coordinate system.

To obtain a representation suitable for statistical analysis, the translation and orientation components were decomposed into interpretable geometric elements and mapped from their native manifolds to Euclidean tangent coordinates.

#### ****3.1 Fréchet means and tangent-space reference****

Fréchet (Karcher) means on $S^{2}$and $SO(3)$ were used to define reference directions and orientations for tangent-space mapping. For a set of points $\left\{ x_{i} \right\}$on a manifold $\mathcal{M}$, the Fréchet mean $\mu\mathcal{\in M}$ is defined as

$$\mu=\arg\min_{v\mathcal{\in M}}\sum_{i} d_{\mathcal{M}}(x_{i},v)^{2},$$

where $d_{\mathcal{M}}$denotes the geodesic distance.

Because this optimization has no closed-form solution in general, means were computed using **Karcher iteration**^1,2^. Starting from an initial estimate, the procedure iteratively:

1. maps each point to the tangent space at the current estimate using the logarithm map,
2. averages the resulting tangent vectors,
3. updates the estimate by mapping the mean tangent vector back to the manifold using the exponential map,

until convergence.

We Initialization $S^{2}$ with normalized Euclidean mean and $SO(3)$ with quaternion-averaged mean. Convergence was defined when the norm of the mean tangent vector fell below $1\times{10}^{-11}$ for $S^{2}$ and $1\times{10}^{-13}$ for $SO(3)$, or after a fixed maximum number of iterations. These Fréchet means define the **reference points about which all tangent coordinates are constructed**.

#### 3.1 Translation decomposition: reach and offset direction

The relative translation vector $t_{rel}$ was decomposed into a scalar reach and a unit offset direction:

$$d_{body}=\left\| t_{rel} \right\|, \hat{u}_{body}=\frac{t_{\mathrm{rel}}}{\left\| t_{rel} \right\|},$$

where $d_{body}\in\mathbb{R}^{+}$ represents the distance from the pMHC frame origin to the body-frame origin, and $\hat{u}_{body}\in S^{2}$ represents the direction of displacement on the unit sphere.

This decomposition separates radial displacement (reach) from angular placement over the pMHC surface (offset direction).

#### 3.2 Mapping of offset directions (S² tangent coordinates)

Offset directions lie on the unit sphere $S^{2}$⊂ $\mathbb{R}$^3^. To obtain unconstrained coordinates, each direction was mapped to the tangent space at a cohort-level Fréchet mean direction $\mu\in S^{2}$.

The Fréchet mean was computed by iterative Karcher iteration: directions were iteratively projected to the tangent space at the current mean via the logarithm map, averaged, and mapped back using the exponential map until convergence.

The logarithm map at $\mu$ provides a tangent vector in the plane $T_{\mu}S^{2}$, whose norm corresponds to the geodesic angle between $\mu$ and $\hat{u}_{body}$. Each tangent vector was expressed in a two-dimensional orthonormal basis $(e_{1},e_{2})$ defined in the tangent plane.

To ensure interpretability, the tangent basis was anchored to the pMHC groove frame. The first basis direction $e_{1}$ was defined by projecting the groove long axis into the tangent plane, and the second basis direction $e_{2}$ completed the orthonormal basis within the plane. The resulting coordinates, $\mathrm{dir}_{t1}$ and $\mathrm{dir}_{t2}$, describe directional displacement along and across the groove.

The antipodal case ($\hat{u}=-\mu$), where the log map is undefined, did not occur in the cohort because all offset directions remained within a local neighborhood of their respective mean directions.

The basis was anchored to the pMHC groove frame: $e_{1}$ was chosen as the groove long axis projected into the tangent plane at $\mu$, and $e_{2}$ completed the tangent basis within $T_{\mu}S^{2}$. For each unit direction $u$,

$$\log_{\mu}\left( u \right)=\arccos\left( \left\langle\mu,u \right\rangle\right)\cdot\frac{u-\left\langle\mu,u \right\rangle\cdot\mu}{\left\| u-\left\langle\mu,u \right\rangle\cdot\mu\right\|}$$

This tangent vector was expressed in the $\left( e_{1},e_{2} \right)$ basis to yield two offset tangent coordinates, $dir_{t1}$ and $dir_{t2}$. The tangent-vector norm $\left\| \log_{\mu}\left( u \right) \right\|$ equals the geodesic angle between $\mu$ and $u$ on $S^{2}$, in radians.

When $u=\mu$, the tangent vector is defined as zero. The antipodal case $u=-\mu$, for which the log map is undefined, did not occur in the analyzed cohort because offset directions remained far from the antipode of their body-specific mean directions.

#### 3.3 Orientation representation and SO(3) tangent mapping

Relative orientations $R_{\mathrm{rel}}$ were represented using unit quaternions

$$q_{body}= \left( q_{w}, q_{x}, q_{y}, q_{z} \right),$$

which provide a compact and numerically stable representation of rotations in $SO(3)$. Because $q$ and $-q$ represent the same rotation, quaternion signs were canonicalized by enforcing $q_{w}\geq0$, ensuring a consistent hemisphere representation.

Fréchet means on $SO(3)$ were computed analogously to the $S^{2}$ case using Karcher iteration. At each iteration, quaternions were mapped to the tangent space at the current mean, averaged, and mapped back via the exponential map until convergence.

To obtain tangent coordinates, relative orientations were mapped using the logarithm map on $SO(3)$. Differences were computed using the right-translated convention,

$$q_{\mathrm{diff}}=q_{\mathrm{body}}\cdot\bar{\mu}_{q},$$

where $\bar{\mu}_{q}$ denotes the quaternion conjugate of the mean. The resulting quaternion difference was converted to an axis–angle form, yielding a tangent vector whose direction encodes the rotation axis and whose magnitude encodes the geodesic rotation angle.

The tangent vector was expressed in the pMHC groove frame, yielding three orientation coordinates $\mathrm{rot}_{t1}$, $\mathrm{rot}_{t2}$, $\mathrm{rot}_{t3}$ corresponding to rotations about:

- $\mathrm{rot}_{t1}$: the groove long axis (roll)
- $\mathrm{rot}_{t2}$: the in-groove perpendicular axis (pitch)
- $\mathrm{rot}_{t3}$: the groove-normal axis (twist)

The use of the right-translated convention ensures that orientation differences are expressed in a common pMHC-aligned coordinate system, enabling direct biological interpretation across structures. Left-translated alternatives would instead express differences in the body frame and were not used here.

#### 3.4 Combined tangent representation

Each body contributes:

- one reach coordinate
- two offset-direction tangent coordinates
- three orientation tangent coordinates

yielding six Euclidean coordinates per body. Across the three bodies (whole TCR, CDR3α, CDR3β), this produces an 18-dimensional tangent representation for each complex.

This representation enables coordinate-level statistical analyses while preserving the geometric meaning of the original manifold-valued descriptors.

#### 3.5 Tangent-space validity and spread diagnostics

The validity of the tangent-space approximation depends on the spread of data around the Fréchet mean. To assess this, geodesic distances from the mean were recorded:

- angular deviations for $S^{2}$offset directions
- geodesic rotation distances for $SO(3)$ orientations

Across the full cohort ($n=378$), all blocks exhibited distributions confined to local neighborhoods of their respective means, ensuring that tangent-space linearization was appropriate.

Additional diagnostics confirmed representation stability, including convergence of Karcher iterations and consistency of mapped coordinates under small perturbations. These checks verify that the Euclidean tangent representation provides a faithful approximation to the underlying manifold geometry for downstream analyses.

### Supplementary Note 4. Statistical analyses for descriptor distributions

This section describes statistical analyses used to characterize FramePose descriptor distributions, compare classes, and relate FramePose variables to conventional descriptors.

#### 4.1 Native manifold class-mean comparisons.

Class-associated geometric differences were first evaluated directly in native descriptor space, without tangent projection.

For each FramePose block, class-specific means were computed using geometry-appropriate definitions:

- Reach ($\mathbb{R}^{+}$): arithmetic mean (Å)
- Offset direction ($S^{2}$): Fréchet (Karcher) mean direction
- Orientation ($SO(3)$): Fréchet mean quaternion

Class shifts were defined as:

- absolute difference (reach, Å)
- geodesic angle on $S^{2}$(offset, degrees)
- quaternion geodesic distance on $SO(3)$(degrees)

Conventional descriptors were treated analogously:

- TCR-CoM ($\theta,\phi$) → embedded as unit vectors on $S^{2}$
- Crossing angle → scalar angular difference

These native manifold class-shift summaries are reported in Table 1.

#### 4.2 Shared tangent-space class tests.

To test class differences in a unified coordinate system, all descriptors were expressed in tangent space at pooled-cohort Fréchet means.

- **Scalar features** (reach, crossing angle):
  → two-sided Mann–Whitney U test
  → effect size:

$$r^{2}=\frac{Z^{2}}{N}$$

- **Multicomponent features** (offset, orientation, TCR-CoM):
  → Hotelling’s $T^{2}$test in tangent coordinates

The $T^{2}$statistic was converted to an $F$-statistic with appropriate degrees of freedom.

$$\text{pseudo-}R^{2}=\frac{T^{2}}{T^{2}+n-2}$$

All p-values were adjusted using Benjamini–Hochberg false discovery rate (FDR) correction.

#### 4.3 Principal-component analysis of FramePose tangent coordinates.

Principal component analysis (PCA) was applied to the 18-dimensional FramePose tangent representation.

To ensure balanced contributions, each FramePose block (reach, offset, orientation) was centered and scaled by its root-mean-square per-coordinate standard deviation

This avoids over-weighting multicomponent features (e.g., SO(3) orientation) the following were recorded:

- explained variance ratios
- principal component scores
- coordinate loadings

For interpretability, block contributions were computed as:

$$\text{block contribution}=\sum(\text{loadings}^{2})$$

summed over coordinates within each FramePose block.

#### 4.4 Native FramePose block dependence by distance correlation.

Dependence between FramePose blocks was quantified using distance correlation applied to native manifold distances.

For each block distance is defined as:

- **Reach**:

$$d_{ij}=\mid d_{i}-d_{j}\mid$$

- **Offset direction** (S²):

$$d_{ij}=\arccos(u_{i}\cdot u_{j})$$

- **Orientation** (SO(3)):

$$d_{ij}=2\arccos(\mid q_{i}\cdot q_{j}\mid)$$

Given double-centered distance matrices $A$, $B$:

$$dCor^{2}\left( A,B \right)=\frac{1}{n^{2}}\sum_{i,j} A_{ij}B_{ij},$$

$$dCor\left( A,B \right)=\frac{dCov\left( A,B \right)}{\sqrt{dCov\left( A,A \right)dCov\left( B,B \right)}}.$$

Significance was assessed by 999 permutations of one distance matrix, and Benjamini–Hochberg correction was applied across the 36 unique off-diagonal block pairs. This native, distance-based analysis is basis-invariant and does not require tangent-space projection. The resulting 9 × 9 distance-correlation matrix is reported in Supplementary Figure S2.

### Supplementary Note 5. Modeling, attribution, and descriptor relationships

This note describes the statistical procedures used to evaluate predictive associations between docking geometry and functional outcomes, to localize contributions of FramePose components, and to quantify redundancy between FramePose and conventional docking descriptors.

#### 5.1 Outcomes and cohort definition

Buried surface area (BSA, Å²) was obtained from TCR3d metadata and analyzed as a continuous outcome. FramePose-based BSA models were fit on the full FramePose cohort (n = 378), and conventional-comparison BSA models were fit on the subset with complete conventional descriptors (n = 377).

Binding affinity was analyzed using curated $K_{D}$ values reported in μM. Entries annotated with “n.d.” (no measurement) were excluded. Entries reported as censored lower bounds (e.g., >X μM) were retained using the reported bound value as the censoring value. This yielded a final affinity cohort of n = 244 complexes for affinity association analyses.

For binary affinity-label analyses, $K_{D}$ values were binarized at 20 μM, with complexes assigned as strong binders when $K_{D}<20$μM (n = 150) and weak binders when $K_{D}\geq20$μM (n = 94). The 20 μM threshold was selected within the 1–100 μM range because it provided a relatively balanced class composition for stable cross-validated association analysis.

Because related receptors, peptides, and MHC alleles may appear in both training and validation folds under random partitioning, affinity analyses are interpreted as cross-validated association analyses within the available cohort rather than deployable affinity-prediction models.

#### 5.2 Cross-validated association modeling

Association modeling was performed for two tasks: (i) BSA regression and (ii) binding affinity classification. All models were implemented using gradient-boosted decision trees (HistGradientBoostingRegressor and HistGradientBoostingClassifier).

Model performance was evaluated using repeated five-fold cross-validation with 20 repeats. Regression used standard K-fold splitting, while classification used stratified K-fold splitting to preserve class proportions. For each repeat, predictions from validation folds were aggregated and performance metrics were computed from pooled out-of-fold (OOF) predictions.

Association strength was quantified using coefficient of determination ($R^{2}$) and Spearman rank correlation for regression, and AUROC and AUPRC for classification. The AUROC chance level is 0.5, and the AUPRC chance level corresponds to the positive-class fraction (150/244 = 0.615 in this cohort).

All models used fixed hyperparameters: maximum iterations = 200, learning rate = 0.05, maximum depth = 4, minimum samples per leaf = 10, and L2 regularization = 1.0.

FramePose features were evaluated using hierarchical configurations (single-body, two-body, and all-body). For each task, the saturated FramePose model was defined as the configuration achieving the highest cross-validated association strength. For BSA regression, this corresponded to the all-body model, whereas for affinity classification, the highest-performing configuration consisted of the CDR3α and CDR3β bodies.

Conventional descriptor models were evaluated using TCR-CoM coordinates, crossing and incident angles, TCRdock parameters, and the combined 16-feature descriptor set. This combined descriptor set served as the baseline model for augmentation analyses.

#### 5.3 Feature attribution analyses

Feature attribution analyses were performed on the task-specific saturated FramePose model to quantify the contribution of geometric components.

In leave-one-block-out (LOO) analysis, model performance was recomputed after removing each FramePose block (reach, offset, or orientation for each body). The change in performance was defined as

$$\Delta=\mathrm{Performance}_{\text{full}}-\mathrm{Performance}_{-\text{block}},$$

which measures the dependence of the fitted model on that block.

Permutation importance was computed by randomly permuting feature values within each block in the validation data while preserving marginal distributions. The resulting change in performance,

$$\Delta=Performance_{\mathrm{original}}-Performance_{\mathrm{permuted}},$$

quantifies the association signal encoded by that block.

Together, LOO and permutation analyses distinguish model dependence from signal strength within the data.

#### 5.4 Descriptor recoverability

Recoverability analyses were performed to assess how much information in FramePose descriptors can be reconstructed from conventional docking descriptors. All analyses used the dataset with complete conventional descriptors (n = 377).

FramePose was represented by 18 tangent coordinates, consisting of reach (scalar), two offset-direction tangent coordinates, and three orientation tangent coordinates for each of the whole-TCR, CDR3α, and CDR3β frames. For block-level analyses, these coordinates were grouped into nine blocks defined by body and feature type (reach, offset, orientation).

Conventional descriptors were represented by a 16-dimensional feature vector. Scalar descriptors (e.g., TCR-CoM radius, TCRdock distance) were used directly. Circular descriptors (TCR-CoM $\theta$ and $\phi$, TCRdock torsion, crossing angle, incident angle) were represented using cosine–sine embeddings. Directional descriptors were expressed as $S^{2}$ tangent coordinates at cohort-level Fréchet means.

FramePose features (coordinates or blocks) were regressed on the 16-dimensional conventional descriptor vector. Scalar targets were evaluated using out-of-fold $R^{2}$. Multicomponent targets were evaluated using joint $R^{2}$, computed by aggregating residual and total sums of squares across target components. For $S^{2}$ and $SO(3)$ blocks, errors were additionally summarized as mean tangent-space residual magnitudes (degrees).

Block-level recoverability was reported for all nine FramePose blocks, and per-coordinate recoverability was reported for each of the 18 tangent coordinates. Empirical 95% intervals were estimated from the distribution of repeated cross-validation scores.

High recoverability indicates that a FramePose feature can be reconstructed from conventional descriptors, whereas low recoverability indicates geometric variation not captured by conventional descriptor families.

To further assess redundancy between descriptor families, we performed reverse-incremental analysis. In this framework, models were first trained using the saturated FramePose feature set for the task of interest. Conventional descriptor sets were then added on top of the saturated FramePose model, and the change in cross-validated association strength was evaluated as

$$\Delta=Performance_{\mathrm{augmented}}-Performance_{\mathrm{FramePose}},$$

This analysis tests whether conventional descriptors provide additional association signal beyond that already captured by FramePose. Minimal or non-significant gains indicate that the corresponding conventional descriptors are largely redundant with the FramePose representation.

#### 5.5 Augmentation analysis relative to conventional descriptors

To evaluate whether FramePose features provide information beyond conventional descriptors, augmentation analysis was performed.

A baseline model was first trained using the combined 16-feature conventional descriptor set. A second model was then trained using both the conventional descriptors and selected FramePose features. The change in performance was defined as

$$\Delta=Performance_{\mathrm{augmented}}-Performance_{\mathrm{baseline}},$$

where performance corresponds to $R^{2}$ for regression and AUROC/AUPRC for classification.

Augmentation analyses used the same repeated five-fold cross-validation scheme as the primary association models, ensuring consistency of evaluation. Positive augmentation gain indicates that FramePose features capture association signals not contained in conventional descriptors.

#### 5.6 Relationship between recoverability and augmentation

To characterize redundancy between descriptor families, the relationship between descriptor recoverability and augmentation performance was examined at the level of FramePose blocks.

Each block was summarized by its recoverability from conventional descriptors, measured as out-of-fold $R^{2}$, and by its augmentation gain when added to the conventional baseline model. Associations between these quantities were quantified using both Spearman and Pearson correlation coefficients.

A negative association between recoverability and augmentation gain indicates that features poorly captured by conventional descriptors contribute disproportionately to predictive performance when added to baseline models. This pattern identifies sources of nonredundant geometric information within the FramePose representation.

### Supplementary Note 6 — Section 3 Analytical Framework

#### 6.1 Biological grouping and contrast design

Biological organization of FramePose geometry was assessed using sequence- and antigen-context grouping variables derived from curated TCR3d metadata and IMGT/ANARCI-annotated TCR regions.

Each Section 3 analysis was formulated as a conditioned focal-factor contrast of the form:

$$focal factor | conditioning variables$$

For example, (V-region | MHC allele) tests whether germline V-region identity explains pose variation within MHC alleles, whereas (CDR3 | pMHC + V-region) tests whether CDR3 sequence explains residual variation after both antigen context and germline framework are fixed.

#### 6.2 Feasibility filtering for conditioned contrasts

Not all candidate contrasts are identifiable due to uneven sampling across grouping variables. For each contrast, conditioning strata were retained only if they contained at least two distinct levels of the focal factor, ensuring that focal effects could be estimated within each stratum.

A feasibility analysis recorded, for each contrast:

- number of usable strata
- number of retained structures
- number of replicated focal levels within strata

These summaries were used to define the valid contrasts analyzed in Section 3.

#### 6.3 Native FramePose distance matrices

Partial PERMANOVA was performed primarily on native FramePose distances, rather than on the 18-coordinate tangent matrix.

For each FramePose block, pairwise distances were computed on the appropriate manifold:

- Reach ($\mathbb{R}^{+}$): absolute Euclidean distance
- Offset ($S^{2}$): geodesic distance $d\left( i,j \right)=\arccos\left( u_{i} \cdot u_{j} \right)$
- Orientation ($SO(3)$): quaternion geodesic distance $d\left( i,j \right)=2\arccos\left| \left\langle q_{i}, q_{j} \right\rangle\right|$

These distances were computed for all combinations of body (whole TCR, CDR3α, CDR3β) and feature type (reach, offset, orientation), yielding nine native FramePose block distance matrices.

Composite distances were then constructed:

- Body-level distances: average of reach, offset, and orientation blocks for each body
- Whole-pose distance: mean-normalized average across all nine blocks

Mean normalization ensured comparable contribution from translational and rotational components.

#### 6.4 PERMANOVA statistic

For a dataset of n complexes partitioned into $G$ groups with pairwise distance matrix $D$, PERMANOVA partitions the total distance-based sum of squares as described by Anderson^3^:

$$SS_{total}=\frac{\Sigma_{\left\{ i,j \right\}}D_{ij}^{2}}{2n}$$

$$SS_{within}=\Sigma_{g}[\frac{\Sigma_{\left\{ i,j \in g \right\}D_{ij}^{2}}}{2n_{g}}]$$

$$SS_{between}=SS_{total}-SS_{within}.$$

The pseudo-F statistic was computed as

$$F =\frac{\frac{SS_{between}}{G - 1}}{\frac{SS_{within}}{n - G}} ,$$

And the distance-based effect size was reported as

$$R^{2}=\frac{SS_{between}}{SS_{total}}.$$

Statistical significance was assessed using 999 permutations of group labels (with restriction when applicable), with p-values computed using the standard $(+1)/(n_{perm}+1)$ correction.

#### 6.5 Conditioned partial PERMANOVA

To isolate the contribution of each biological determinant, Type-II partial PERMANOVA was performed under the conditioned contrast design.

Let:

- $R_{cond}^{2}$: variance explained by conditioning variables
- $R_{joint}^{2}$: variance explained by conditioning variables plus focal factor

The focal effect is defined as

$$\Delta R^{2}=R_{joint}^{2}-R_{cond}^{2}.$$

To control for inflation due to high group cardinality, an adjusted $\Delta R^{2}$ was computed by applying adjusted-$R^{2}$ corrections to both models prior to differencing. This adjusted $\Delta R^{2}$ was used as the primary effect size, particularly for high-cardinality groupings such as V-region, receptor identity, and CDR3 sequence.

Statistical significance of $\Delta R^{2}$ was assessed using restricted permutation testing. For each contrast:

- focal labels were permuted only within conditioning strata
- conditioning structure was preserved
- focal association with geometry was disrupted

A total of 999 restricted permutations were performed for each test. Permutation p-values were computed with the same continuity correction described above. Benjamini–Hochberg false discovery rate correction was applied within each contrast family across tested blocks and axes.

#### 6.6 Block-level and axis-level decomposition

Primary tests were performed on the whole-pose composite distance to assess overall determinant effects.

To localize effects:

- Block-level PERMANOVA was applied to the nine native FramePose blocks
- Axis-level PERMANOVA was applied to the 18 tangent coordinates

For axis-level analysis, one-dimensional Euclidean distance matrices were constructed from absolute coordinate differences. The same conditioned partial PERMANOVA framework was applied.

- Block-level analyses determine whether effects localize to reach, offset, or orientation in specific bodies
- Axis-level analyses identify specific geometric directions (e.g., groove-normal twist, cross-groove pitch)

#### 6.7 Recrystallization reproducibility floor and pairwise distance scale

To provide a physical scale for interpreting native whole-pose distances, repeated-complex groups were audited using recryst_floor_audit.py. Candidate repeated complexes were initially defined by shared receptor identity and shared peptide–MHC context. Each multi-member group was then audited to determine whether it represented a true recrystallization rather than a distinct engineered construct. Groups were retained as true recrystallizations only if receptor identity, peptide sequence, MHC chain sequence, and mutation status were identical, and if the peptide annotation was unambiguous. Groups containing ambiguous non-natural peptide residues, engineered MHC mutations, altered TCR mutations, or different MHC-chain sequences were excluded from the reproducibility-floor estimate.

Native whole-pose composite distances were computed among structures within the audited true-recrystallization groups. The pooled median of these pairwise distances was used as the empirical FramePose reproducibility floor. This floor was then used to contextualize broader pairwise distance categories, including same receptor and same peptide–MHC, same germline V-region with different CDR3, same germline V-region with different peptide–MHC context, different germline V-region within the same peptide–MHC, and all different-germline pairs. These pairwise summaries were used as a distance-scale interpretation of the partial-PERMANOVA results.

#### 6.8 PERMDISP dispersion diagnostic.

For selected PERMANOVA contrasts, we additionally tested whether significant group structure could be explained by unequal within-group dispersion rather than differences in group location. PERMDISP was applied to the same native FramePose distance matrices used for PERMANOVA. For each grouping variable, the distance matrix was first principal-coordinate decomposed, and each complex was represented in the resulting Euclidean principal-coordinate space. Within each group, the multivariate centroid was computed, and each complex’s distance to its group centroid was recorded. The PERMDISP statistic tests whether the mean distance to centroid differs among groups.

For a grouping with $G$ groups, let $z_{ig}$ denote the distance of complex $i$ in group $g$ to that group’s centroid. Group dispersions were summarized as the mean centroid distance,

$$\bar{z}_{g}=\frac{1}{n_{g}}\sum_{i\in g} z_{ig}.$$

A one-way ANOVA was performed on centroid distances, and significance was assessed using 999 permutations, with restrictions matching the corresponding PERMANOVA design.

- Non-significant PERMDISP: supports interpretation of PERMANOVA results as location effects
- Significant PERMDISP: indicates possible contribution of dispersion differences and was interpreted cautiously

### Supplementary Note 7. CDR3-epitope contact-count definitions

#### For each structure, we removed hydrogen and deuterium atoms, and retained heavy atoms from TCR, peptide, and MHC residues. TCR$\alpha$ and TCR$\beta$ chains and CDR regions were assigned either by IMGT/ANARCI numbering, when available, or by metadata-defined CDR spans. Contact summaries used CDR3$\alpha$ and CDR3$\beta$ residues as TCR-side regions and peptide or peptide+MHC (pMHC) residues as pMHC-side targets.

#### Candidate TCR–pMHC residue pairs were prefiltered by C$\alpha$–C$\alpha$ distance. A residue pair was evaluated if both residues had C$\alpha$ atoms within 16 Å, a permissive prefilter chosen to avoid missing long side-chain contacts from residues such as Arg, Lys, or Trp. For each retained residue pair, all heavy-atom pairs were evaluated. The master heavy-atom contact cutoff was 4.0 Å by default, and this cutoff was recorded in the output as contact_cut. Sensitivity runs could be performed with alternative cutoffs, such as 5.0 Å.

#### For each residue pair, the script reported both atom-pair event counts and residue-pair presence indicators. Total contacts (n_contacts) were defined as the number of heavy-atom pairs within the master cutoff. Salt bridges were defined as oppositely charged atom pairs within the salt-bridge cutoff of 4.0 Å. Acidic oxygen atoms from Asp/Glu side chains and C-terminal OXT atoms were treated as anions, and Arg/Lys/His nitrogen atoms were treated as cations. The salt-bridge event count was reported as (n_salt_bridges).

#### Hydrogen bonds were defined using donor–acceptor atom-pair geometry. Backbone nitrogen atoms, except in Pro, and annotated side-chain donor atoms were treated as donors when the corresponding antecedent atom was present. Backbone oxygen atoms and annotated side-chain acceptor atoms were treated as acceptors. A hydrogen bond required donor–acceptor distance ≤3.5 Å and an antecedent–donor–acceptor angle ≥120°. By default, charged hydrogen bonds were assigned to the salt-bridge class rather than double-counted as hydrogen bonds; the script also allowed an independent classes option in which a charged hydrogen bond could be counted in both classes. Hydrogen-bond events were reported as (n_hbonds).

#### Polar contacts were defined as the sum of salt-bridge and hydrogen-bond events. The event count was reported as (n_polar).

#### Packing was summarized in two ways. First, legacy atom-partition counts were reported for reference: n_hydrophobic counted apolar C/S–C/S atom pairs within the master contact cutoff after excluding salt-bridge and hydrogen-bond atom pairs, and n_vdw counted the remaining nonpolar/nonclassified atom contacts. Methionine sulfur (SD) and cysteine sulfur (SG) were treated as apolar by default to include sulfur-mediated packing. Second, to reduce residue-size bias from large side chains, the primary packing descriptor was a continuous residue-pair packing score, w_pack, computed from the minimum heavy-atom distance of the residue pair using a logistic switching function,

##

$$w_{pack}=\frac{1}{1+exp\{k(d_{min}-d_{0})\}},$$

#### with default midpoint $d_{0}$=4.0 Å and steepness $k$=4.0. Residue pairs with no atom pair within the master contact cutoff were still emitted if their minimum heavy-atom distance fell within the soft packing window of 6.0 Å. The per-summary packing score, $S_{pack}$, was defined as the sum of $w_{pack}$ over residue pairs in a given loop–target partition.

1.	Karcher H. Riemannian center of mass and mollifier smoothing. *Commun Pure Appl Math*. 1977;30(5):509-541. doi:10.1002/cpa.3160300502

2.	Pennec X. Intrinsic Statistics on Riemannian Manifolds: Basic Tools for Geometric Measurements. *J Math Imaging Vis*. 2006;25(1):127-154. doi:10.1007/s10851-006-6228-4

3.	Anderson MJ. A new method for non-parametric multivariate analysis of variance. *Austral Ecol*. 2001;26(1):32-46. doi:10.1111/j.1442-9993.2001.01070.pp.x
